# The role of anatomical connection strength for interareal communication in macaque cortex

**DOI:** 10.1101/2020.12.15.422902

**Authors:** Julien Vezoli, Martin Vinck, Conrado Arturo Bosman, André Moraes Bastos, Christopher Murphy Lewis, Henry Kennedy, Pascal Fries

## Abstract

What is the relationship between anatomical connection strength and rhythmic synchronization? Simultaneous recordings of 15 cortical areas in two macaque monkeys show that interareal networks are functionally organized in spatially distinct modules with specific synchronization frequencies, i.e. frequency-specific functional connectomes. We relate the functional interactions between 91 area pairs to their anatomical connection strength defined in a separate cohort of twenty six subjects. This reveals that anatomical connection strength predicts rhythmic synchronization and vice-versa, in a manner that is specific for frequency bands and for the feedforward versus feedback direction, even if interareal distances are taken into account. These results further our understanding of structure-function relationships in large-scale networks covering different modality-specific brain regions and provide strong constraints on mechanistic models of brain function. Because this approach can be adapted to non-invasive techniques, it promises to open new perspectives on the functional organization of the human brain.

## INTRODUCTION

The quest to understand integrated brain function has motivated a shift in emphasis away from single neurons or brain areas and towards circuits and large-scale networks (Markov et al., 2013; Rubinov and Sporns, 2011). While considerable theoretical and modelling work has treated neuronal networks as random graphs (Maass et al., 2002; Morrison et al., 2007), recent network analyses have introduced an exponential distance rule (EDR) based on the dependence of anatomical connection strength with distance. The EDR explains numerous features of interareal anatomical connectivity and together with work at the local circuit level (Cossell et al., 2015; Ko et al., 2011; Ohki et al., 2005) suggests that brain organization is not random, but highly structured with characteristic features that depend centrally on the distribution of connection weights (Ercsey-Ravasz et al., 2013; Horvát et al., 2016; Theodoni et al., 2020). Anatomical connectivity provides the backbone for functional interactions, for example, as estimated by inter-areal correlations. Brain-wide activity measurements using functional magnetic resonance imaging (fMRI) led to the discovery of highly structured brain-wide networks (Bassett and Bullmore, 2009; Biswal, 2012; Fox et al., 2005). Those networks partly correlate with patterns of anatomical connectivity, and they predominantly reflect ongoing, intrinsic activity rather than sensory inputs (Fox et al., 2005; Honey et al., 2009). Those functional networks have been explored in topographical (Margulies et al., 2016) and graph-theoretic terms (Bassett and Sporns, 2017), partly relating functional networks to different metrics of anatomical connectivity (Park and Friston, 2013). Importantly, intrinsic functional networks are related to stimulus- and/or task-dependent networks (Cole et al., 2014; Fox et al., 2005; Lewis et al., 2016), and able to influence cognitive functions (Baldassarre et al., 2012; Cohen, 2018; Cole et al., 2016; Lewis et al., 2009).

A candidate mechanism for neuronal interactions in large-scale networks is rhythmic neuronal synchronization (Buzsáki and Draguhn, 2004; Engel et al., 2001; Fries, 2015; Kopell et al., 2014; Lynn and Bassett, 2019; Palmigiano et al., 2017; Salinas and Sejnowski, 2001; Schnitzler and Gross, 2005; Schoffelen and Gross, 2009; Varela et al., 2001). Several studies using electro- or magnetoencephalography (EEG or MEG) have provided important insights into large-scale networks defined by rhythmic synchronization (Hipp et al., 2012; Roux and Uhlhaas, 2014; Siegel et al., 2008; Uhlhaas et al., 2008; Wang et al., 2019). One of the key findings is that networks defined by rhythmic synchronization typically differ markedly between the characteristic frequency bands, such as the theta, alpha and beta bands. Combinations of MEG-based analysis of rhythmic synchronization networks with MRI-based measurements of structural and functional connectivity reveal intriguing links and dissociations (Hipp et al., 2012).

Further, the non-invasive methods available for human subjects are technically limited in terms of exploring structural and functional features of the brain. Activity from different neuronal sources is mixed in the EEG or MEG signals, and unmixing can never be perfect. This is particularly limiting for the study of rhythmic synchronization, which requires measurements of the putatively synchronized brain regions to be independent in order to avoid spurious correlations (Bastos and Schoffelen, 2015; Palva et al., 2018; Schoffelen et al., 2008). Similarly, MRI-based estimates of anatomical connectivity, such as diffusion-tensor-imaging (DTI) based tractography, is only modestly correlated with ground-truth anatomical data (Donahue et al., 2016). These limitations can in principle be overcome in animal models that allow invasive electrophysiological recordings and tracer-based anatomy.

Here, we capitalized on two unique datasets, obtained through large-scale electrocorticography (ECoG) recordings and retrograde tract tracing in macaque monkeys. We used 252-channel ECoGs to record local field potentials from large parts of the surface of the left hemisphere of two macaques performing a selective visual attention task. Electrodes covered 15 brain areas, allowing us to investigate different metrics of rhythmic synchronization between 105 pairs of areas. We used data from retrograde tracer injections in 14 areas of 26 macaques, with precise counts of monosynaptically connected neurons, for 91 pairs of areas, separately for supra- and infragranular laminar compartments. Our analysis reveals and characterizes fine-grained spatial correlation structures of large-scale synchronization-defined networks and their specificity for different rhythms. We demonstrate that the strength of cortico-cortical anatomical projections is specifically related to synchronization in the gamma and high-beta bands, but less so in the beta band. The structure-function relationship in the gamma band is primarily with anatomical feedforward projections, and in the beta band with feedback projections. Finally we provide evidence, through overlays of synchronization strengths onto the anatomical core-periphery structure, that the gamma band predominates in the forward periphery whereas lower frequencies are spread across the whole core-periphery structure.

## Results

We investigated neuronal activity in large-scale brain networks in two macaque monkeys performing a selective visual attention task (Fig. 1A, see Methods). Subdural ECoG-grids chronically implanted over large parts of the left hemisphere (Fig. S1), allowed for the simultaneous recording of 15 cortical areas. In the present study, we focused on the task period with sustained visual stimulation and attention. Attention conditions were pooled to increase sensitivity. In order to remove the common recording reference, signals from immediately neighboring electrodes were subtracted from each other to form local bipolar derivations, referred to as (recording) sites.

**Figure 1.**
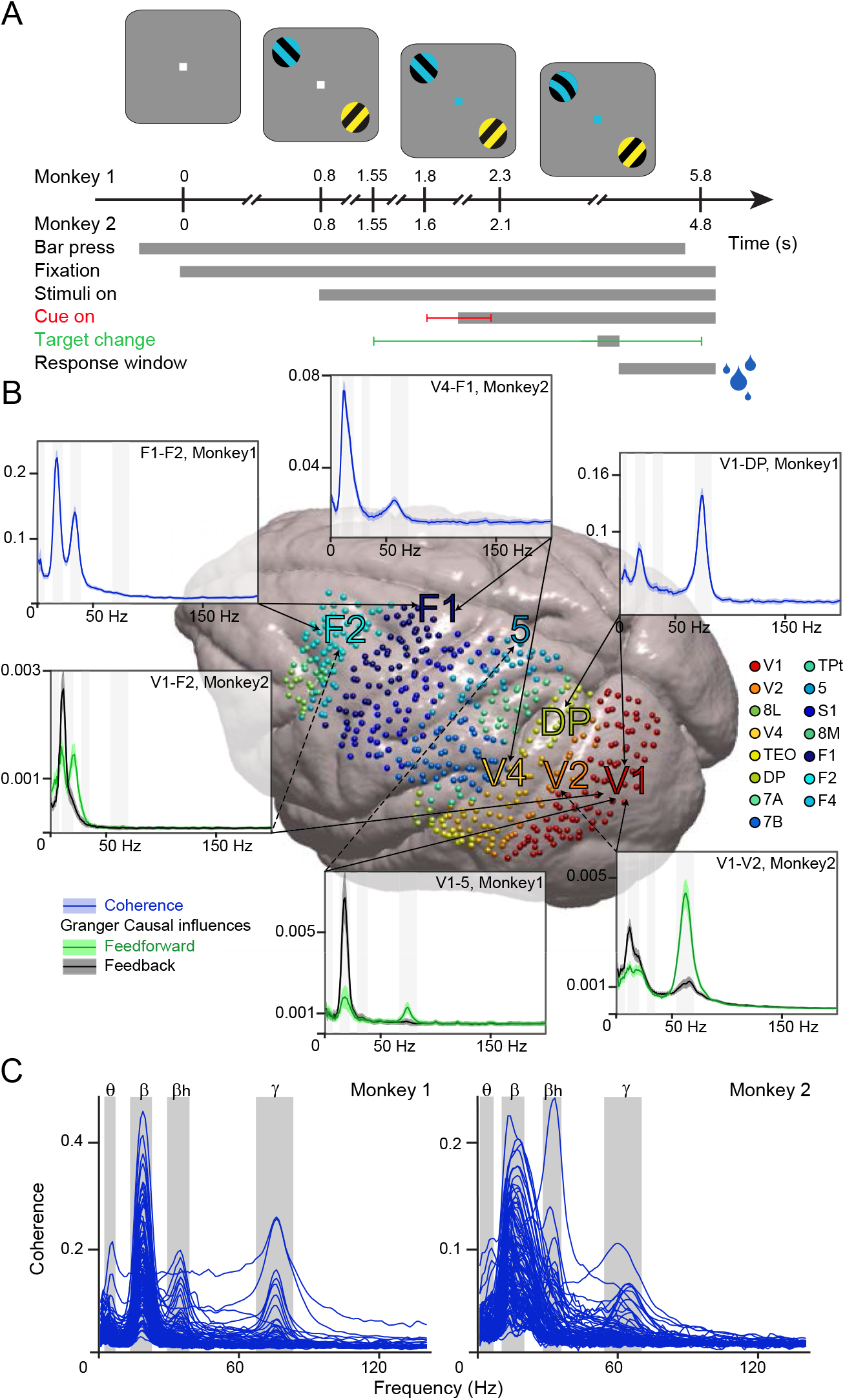
Functional Connectivity datasets recorded during selective visual attention. (A) Two macaque monkeys were trained to release a lever when a change occurred to the peripheral stimulus cued by the color of the fixation dot, while maintaining fixation and ignoring changes to the distractor. Correct performance was rewarded by liquid reward (blue droplets). Task delays for Monkey 1 and 2 in sec. (B) Pooled recording sites of both monkeys on the surface of the INIA19 template brain (see color legend for area assignment). Spectra show examples of interareal Coherence (in blue) and GC (green: feedforward; black: feedback, dashed/plain lines point to the cortical area sending feedback/feedforward projections). Spectra show mean over all trials ±99.9% confidence intervals from bootstrap estimates over trials. (C) Coherence spectra of all 105 area pairs for the two macaque monkeys. Each spectrum represents the mean value of all site pairs for that pair of areas. Shaded areas represent full width at half-maximum (FWHM) for each of the four frequency peaks.

### Interareal functional connectivity (FC) occurs in four characteristic frequency bands

We computed the following frequency-resolved functional connectivity (FC) metrics between all possible site pairs: 1) Coherence, i.e. a metric of interareal synchronization, 2) Power correlation, i.e. the Spearman rank correlation between fluctuations in band-limited power, and 3) Granger causality (GC), i.e. a metric of directed interareal influence. Coherence and power correlation are undirected metrics, whereas GC is a directed metric that allows calculation of spectral influences in both directions. These spectra showed distinct peaks, which are specific for the respective pair of brain areas (see Fig. 1B for example coherence and GC spectra). Across all 105 area pairs, average FC spectra showed peaks for four characteristic brain rhythms, with some individual differences across the two monkeys (Fig. 1C for coherence; Fig. S4A for coherence, power correlation and GC): The theta rhythm (3 ±2Hz in Monkey1 and 4 ±3Hz in Monkey2; peak ±full width at half maximum), the beta rhythm (18 ±5Hz in Monkey1 and 15 ±5Hz in Monkey2), the high-beta rhythm (34 ±5Hz in Monkey1 and 32 ±4Hz in Monkey2) and the gamma rhythm (75 ±8Hz in Monkey1 and 62 ±8Hz in Monkey2) (Fig. 1B-C and S4A).

### Different rhythms show distinct FC network topographies

We first describe spatial patterns for the most prominent spectral bands, the gamma and beta rhythms (Fig. 2). Coherence and power correlation show similar patterns across site pairs within the two frequency bands (Fig. 2A for Monkey 1 and Fig. S2A for Monkey 2; compare upper and lower matrix halves; matrices are provided per monkey, as ECoG coverage and number of recording sites per area differs between monkeys). However, between frequency bands, patterns differed markedly (Fig. 2A, Fig. S2A; compare left and right matrix).

**Figure 2.**
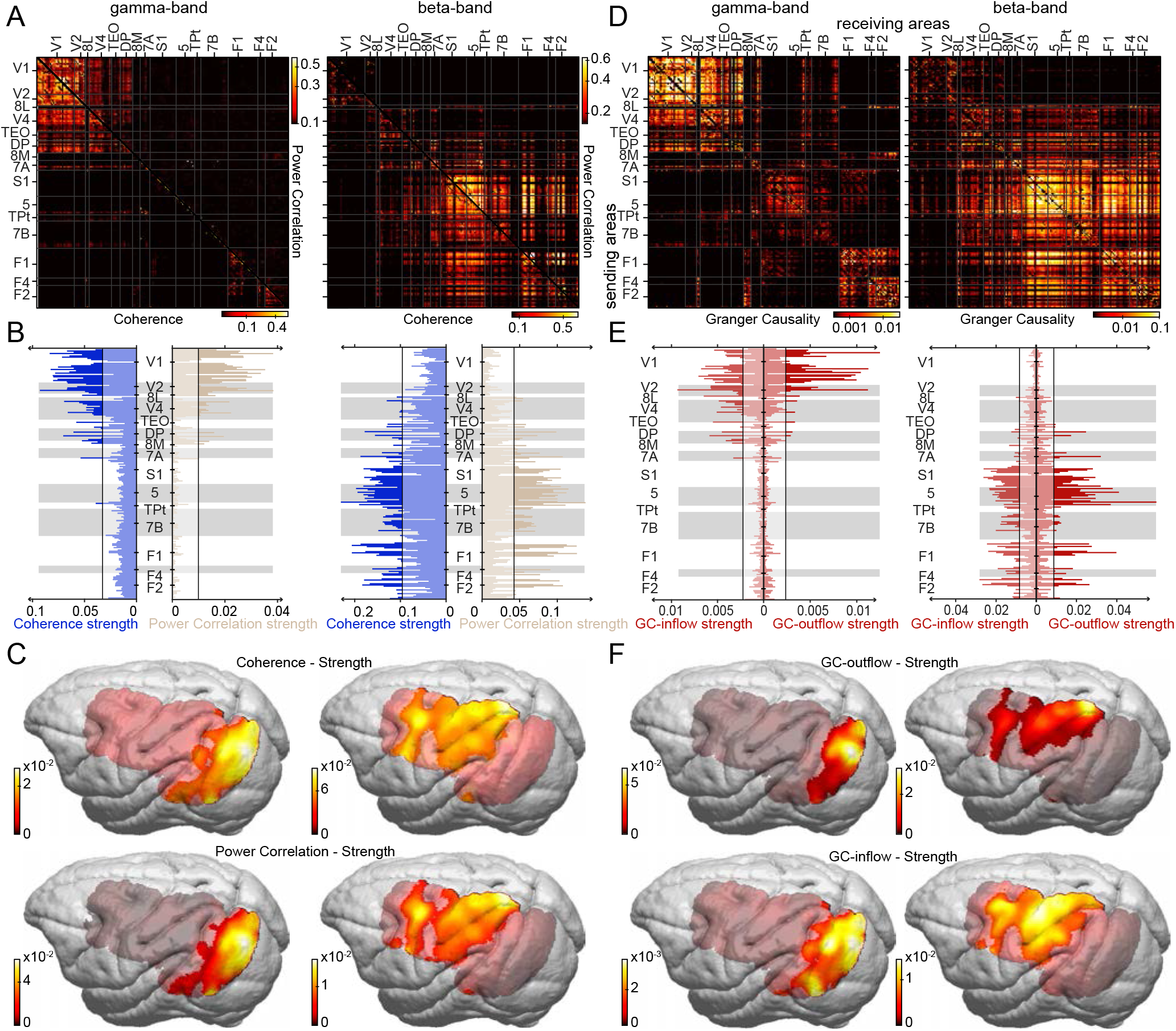
Spatial FC patterns for gamma and beta. (A) The upper parts of the matrices show the power correlations for all possible site pairs of Monkey 1, the lower parts shows the same for coherence (). The axes list the areas, from which the sites had been recorded, with the areas ordered according to their hierarchical level (Chaudhuri et al., 2015). Area boundaries are indicated by gray lines on the matrices. (B) Strength metric for coherence (left) and power correlation (right), for all sites of Monkey 1. The y-axes list the areas, from which the sites have been recorded, ordered as in (A). Area boundaries are indicated by alternating gray shading. (C) Coherence strength (top) and power-correlation strength (bottom) for gamma (left) and beta (right), projected on the template brain and averaged over both monkeys. (D) Similar to (A), but showing the full matrices of GC from all sites of all areas listed on the y-axis (sending areas) to all sites of all areas listed on the x-axis (receiving areas). (E) Similar to (B), but showing the strength metric for GC directed into the respective sites, i.e. GC inflow strength, or out of the respective sites, i.e. GC outflow strength. (F) GC outflow strength (top) and GC inflow strength (bottom) for gamma (left) and beta (right), projected on the template brain and averaged over both monkeys. Note the logarithmic color scales in A and D. Analyses in A-B, D-E shown for Monkey 2 in Fig. S2A-D.

To investigate whether these band-specific patterns of site-pair-wise FC reflect brain topographical patterns, we calculated the strength metric. Coherence strength is defined for each recording site as the average coherence of that site with all other sites (excluding sites within a 2 mm radius to avoid residual volume conduction effects). Power-correlation strength is defined accordingly. Strengths show very similar coherence and power correlation patterns, but very different pattern for gamma versus beta (Fig. 2B for Monkey 1 and Fig. S2B for Monkey 2). Strength projections on the brain reveal that gamma strength peaks in early visual areas (Fig. 2C left; maps are averages over monkeys, after co-registration to a template brain and exclusion of parts covered in only one animal). By contrast, beta strength is highest in the parietal and frontal association areas along the intra-parietal sulcus (IPS) and the precentral gyrus (Fig. 2C right).

GC provides directed FC matrices as shown in Fig. 2D and Fig. S2C for gamma (left) and beta (right). We previously showed that gamma GC between visual areas is typically stronger in the anatomically defined feedforward than in the feedback direction (Bastos et al., 2015). This corresponds to larger values in the upper half–matrix, compared to the corresponding entries in the lower half-matrix, for areas V1 to 7A. Fig. 2D shows that these hierarchy-related differences are superimposed on other spatial patterns: Gamma GC is generally stronger for lower areas, while beta GC is stronger for higher areas, and this overall pattern holds for both inflows and outflows. We calculated the strength metric for GC-inflow and GC-outflow (Fig. 2E and S2D), and the spatial distributions resembled those for coherence and power correlation (Fig. 2F).

The same analyses for the theta and the high-beta band (Fig. 3 and S3A-D) reveal band-specific patterns of site-pair-wise FC and strength topographies that differed from those observed for gamma and beta. The site-pair-wise analysis of theta coherence (Fig. 3A, left) indicates a cluster centered on visual areas V1, V2, V4, TEO, and another centered on motor areas F1, F2, F4, and a substantial distributed FC. The pattern for high-beta is similar to that for theta, but with a clear focus on the motor cluster (Fig. 3A, right). The resulting strength topographies for theta reveals a relatively distributed pattern (Fig. 3C left). By contrast, strength topographies for high-beta reflect the strong focus on motor areas, while excluding prefrontal territory anterior to the arcuate sulcus.

**Figure 3.**
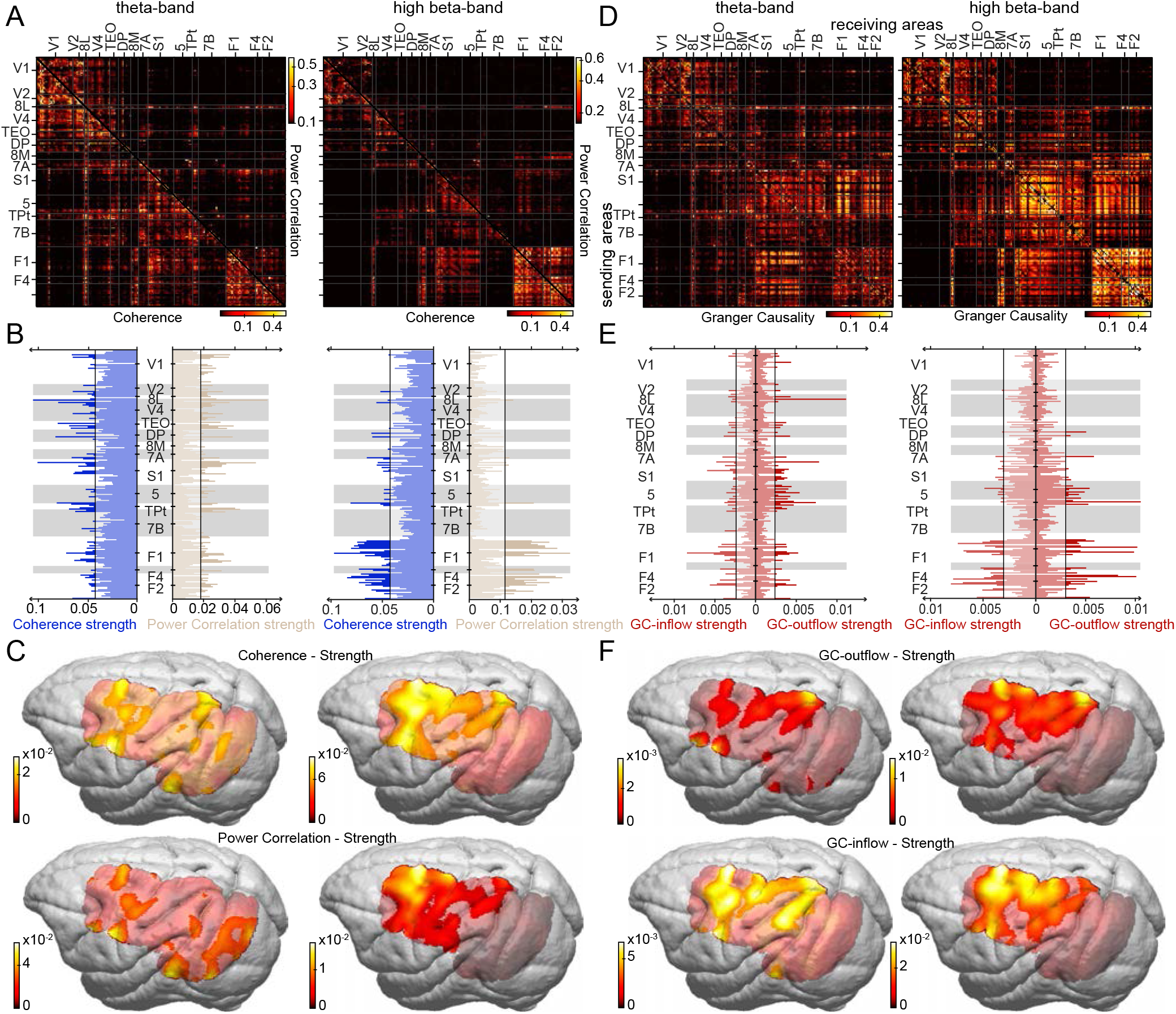
Spatial FC patterns for theta and high-beta. Same as Fig. 2, but for the theta and the high-beta band (Analyses in A-B, D-E shown for Monkey 2 in Fig. S3A-D).

These observations were partly, but not fully, characteristic also for the GC. The full GC matrices (Fig. 3D) resembles the coherence matrices, while even stronger similarities are observed between theta and high-beta. For high-beta, the strength topographies for GC in- and outflow (Fig. 3F, right) strongly resembled those for coherence and power correlation. By contrast, for theta (Fig. 3F, left), the strength topographies for GC in- and outflow were more structured than for coherence and power correlation extending on the gyri either side of the IPS, with a relative sparing of the IPS itself, and with another peak on the precentral gyrus.

We also demonstrate that weak long-distance FC deviates significantly from randomized inter-site FC (see Methods). Significant interareal FC covers long distances (>20mm) for all frequencies; FC at gamma (Fig. S2E) extends up to 50 mm, FC at beta (Fig. S2F), theta and high-beta (Fig. S3E-F) beyond 60 mm.

### FC network topographies correlate with anatomical connectivity

We next investigated whether these FC patterns could be partly explained by known patterns of anatomical connectivity (AC). We previously showed that GC asymmetries are related to the feedforward/feedback character of the respective anatomical projections (Bastos et al., 2015), as quantified by the supragranular labelled neuron (SLN) percentage value (Barone et al., 2000). In the present study we address another fundamental aspect of an anatomical projection namely its strength, which is captured by the fraction of labeled neurons (FLN) (Vezoli et al., 2004). After injection of a retrograde tracer into area A, retrogradely labeled neurons are counted across the brain, e.g. in area B. The FLN of the projection from B to A is the number of labeled neurons found in B divided by the total number of neurons found across the brain. In this way, FLN reflects the fraction of the total synaptic input to A that originates in B. While the majority of projections to a given cortical area arises from within the area itself (~80%), we are interested in projections arising from other areas and so estimate the extrinsically originating fraction of labeled neurons (FLNe). The FLNe excludes counting the neurons inside the injected area, and therefore estimates the fraction of the interareal synaptic input to A that originates in B (Markov et al., 2011).

We were interested in how interareal FC, assessed by coherence, power correlation and GC, relates to interareal AC, assessed by FLNe. We used a dataset based on 28 retrograde tracer injections across 14 cortical areas (Fig. 4A), resulting in a 14X14 matrix of FLNe values (Fig. 4C). Note that this is a directed matrix of AC, in which FLNe from area A to area B is quantified independently of the FLNe in the reverse direction. Thus, the full FLNe matrix can be compared to the full GC matrix. By contrast, the matrices of coherence and power correlation assess overall FC irrespective of direction. To relate them to anatomical connection strength, we averaged FLNe over the two directions, giving a half matrix (Fig. 4B).

**Figure 4.**
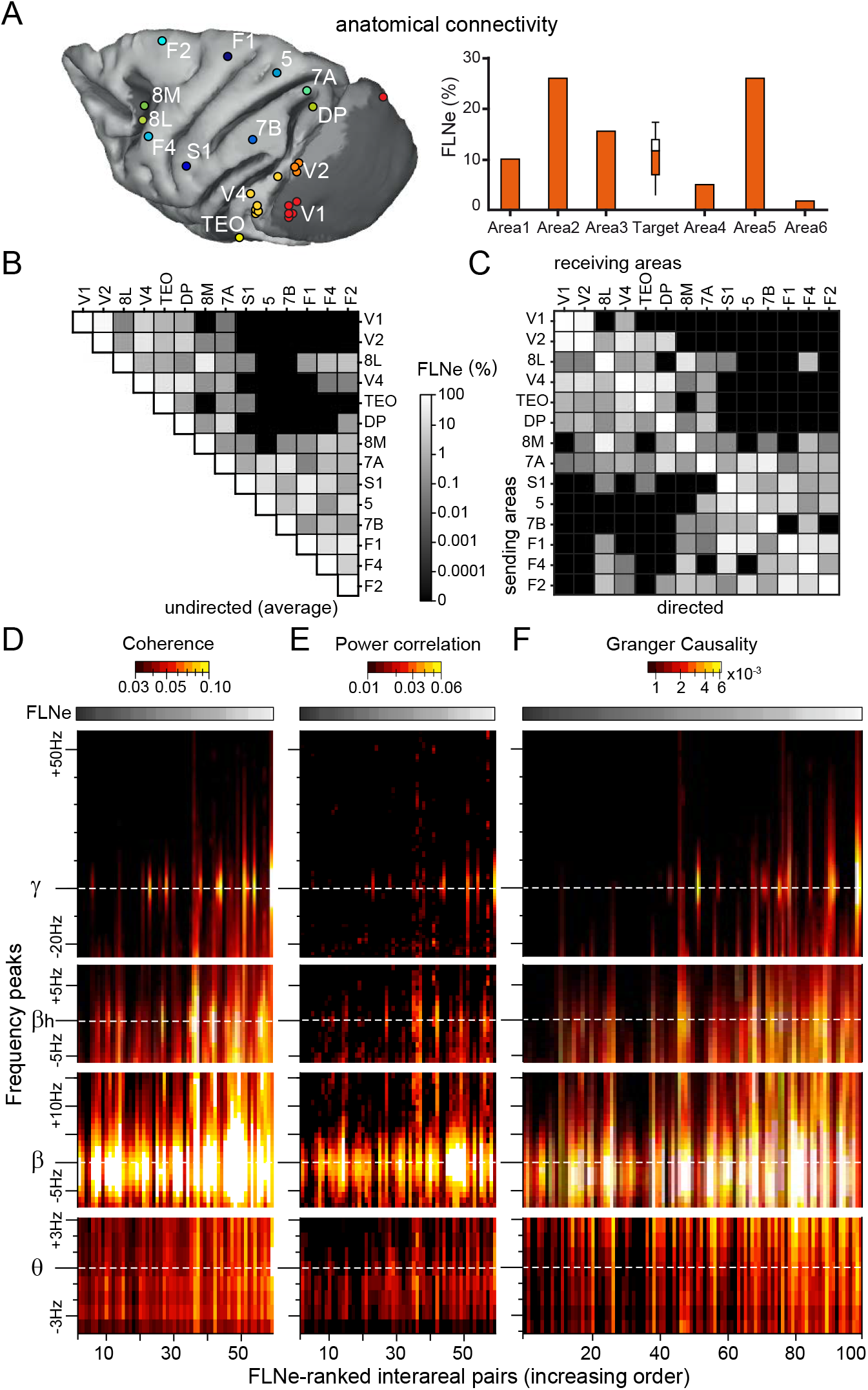
FLNe and FLNe-sorted FC spectra. (A) Each colored dot indicates the injection site of a retrograde tracer, shown here on a template brain. (B) FLNe values for all indicated pairs of areas, averaged over the projections in the respective two directions, e.g. V1-to-V4 and V4-to-V1. Note the logarithmic grey scale, spanning seven orders of magnitude. (C) FLNe values for all indicated projections from the areas listed on the y-axis to the areas listed on the x-axis. Black matrix entries indicate projections for which less than 10 labelled neurons were counted. Those entries were discarded for the average shown in (B). (D) Coherence, power correlation and GC values (color coded) as a function of frequency (y-axis, relative to per-monkey band-wise peak frequencies) and as a function of FLNe rank (x-axis) of the respective interareal pair. For coherence and power correlation, ranking used FLNe averaged over the two directions as shown in (B), and for GC, ranking used FLNe as shown in (C). FC values were first averaged over all respective interareal site pairs and subsequently over animals.

We used this averaged FLNe to rank interareal connections and plot the respective coherence spectra (Fig. 4D). Spectra were first averaged over all site pairs of a given area pair, and subsequently over the two animals. For this, we first determined the peak frequencies per monkey and per rhythm (theta, beta, high-beta, gamma), and expressed frequencies relative to the per-monkey peak frequencies. This revealed that overall, power correlation increased with increasing FLNe. This pattern was also present for power correlation (Fig. 4E) and GC (Fig. 4F).

For this analysis, we excluded FLNe values based on less than 10 labelled neurons, to assure reliability of FLNe estimation (Markov et al., 2014). The pattern held, when we included those FLNe values (Fig. S4B), or when we replaced them by estimates from a model fitted to neuron counts from the non-zero FLNe values (Fig. S4C, see Methods).

### FLNe-FC correlations differ across FC types and across frequencies

To quantify the observed patterns, we performed linear regression analysis between log(FC) and log(FLNe), separately for all combinations of FC type (coherence, power correlation and GC) and frequency band (theta, beta, high-beta, gamma) (Fig. 5). For each combination, there was a significantly positive correlation (P < 4.17E^−03^ after Bonferroni correction for multiple comparisons), yet with a wide range of correlation strengths (Fig. 5A). FLNe was least predictive for FC at beta, with explained variance (R^2^-values) for beta power correlation or beta GC of 0.14. FLNe was most predictive of theta (R^2^=0.47) and high-beta coherence (R^2^=0.39) or gamma GC (R^2^=0.42).

**Figure 5.**
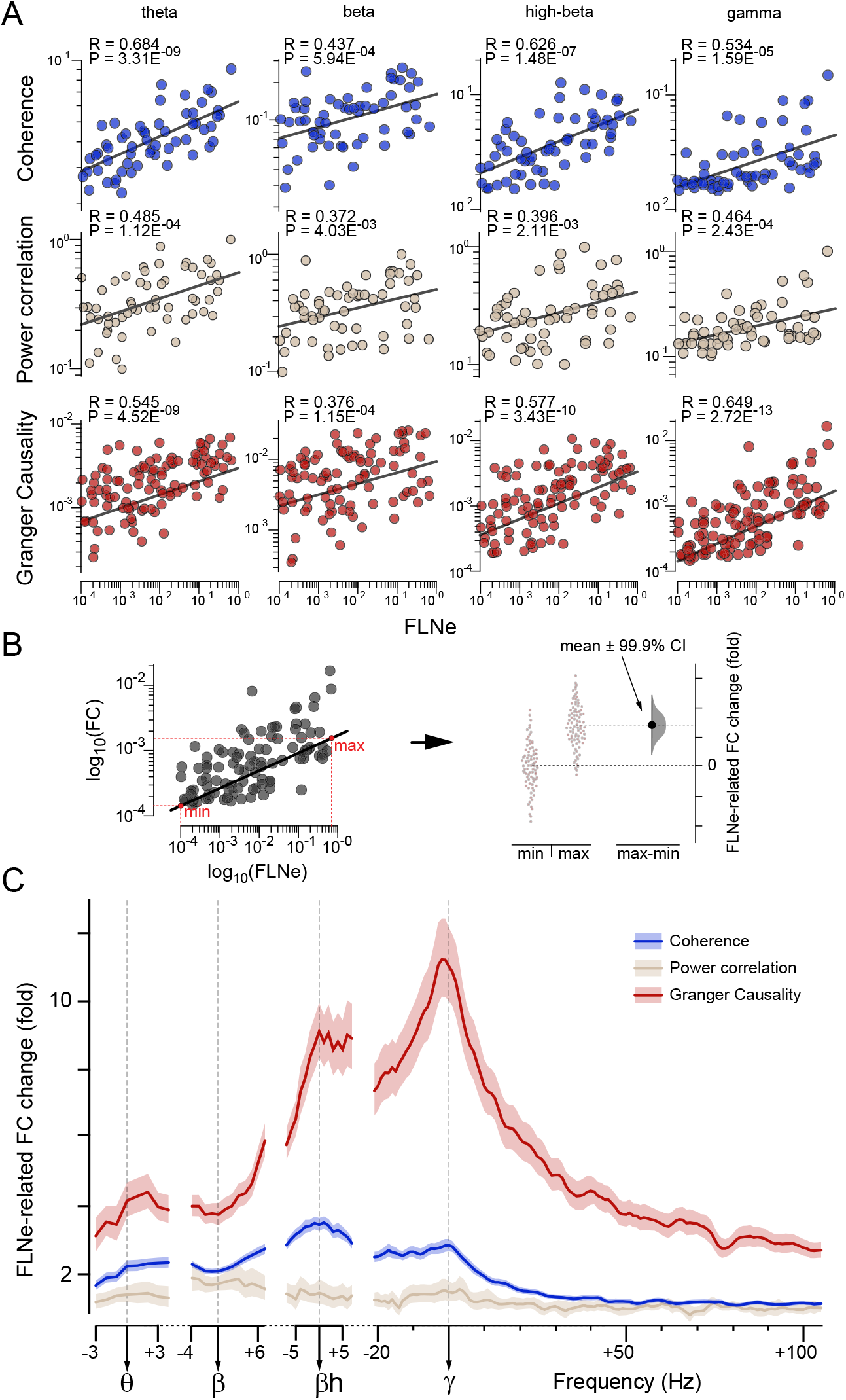
Functional and anatomical connectivity display frequency-dependent covariance. (A) Scatterplots between the three interareal FC measures (from top to bottom: Coherence, Power correlation and Granger Causality) and AC i.e. FLNe. FC values were averaged over monkeys before correlation analysis. (B) With both axes in log10 units, subtraction of FC values between minimum and maximum AC values (left), can be interpreted as FLNe-related FC fold-change (right). (C) FLNe-related FC change as a function of FC frequency. FC spectra (color coded, legend top-right) have been aligned on each frequency-peak before averaging over monkeys (from left to right: theta, beta, high-beta and gamma frequency peaks) and then correlated with AC, i.e. FLNe. Mean over all trials ±99.9% confidence intervals from bootstrap estimates over trials.

To capture the size of the FLNe effect on FC, we used a regression analysis (Fig. 5B). We performed a simple linear regression with the dependent variable FC (i.e. either power correlation, coherence or GC) and the independent variable FLNe. We then used the linear fit to calculate the expected FC at the minimal FLNe value, i.e. FC(min(FLNe)), and at the maximal FLNe value, i.e. FC(max(FLNe)). The ratio FC(max(FLNe)) / FC(min(FLNe)) was used as the FLNe-related FC change (Fig. 5C). This metric is related to the regression slope, but normalizes for differences in FC across frequencies that are not due to FLNe. We derived error estimates by 100 bootstrap replications over trials (Fig. 5B,C) (Efron and Tibshirani, 1994).

This analysis revealed that the three types of FC showed different degrees of dependency from FLNe: Power correlation was least FLNe-dependent, coherence was intermediate, and GC was by far the most strongly FLNe-dependent. The spectrum for power correlation did not show any clear peaks (even when scaled independently). The spectrum for coherence showed peaks for high-beta and gamma, and a local trough for beta, while that for GC showed a small peak in the theta range, and substantial peaks for high-beta and gamma, and again a local trough for beta. These results suggest that the dependence of coherence and GC on AC has a characteristic dependence on frequency: At the individual peak frequencies for gamma and high-beta (and partly also for theta), this dependence is stronger than at neighboring frequencies; by contrast, at the individual peak frequencies for beta, this dependence is weaker than at neighboring frequencies.

### FC-FLNe correlations are not explained by distance, yet FC predicts FLNe

FLNe has been shown to decrease exponentially with interareal distance, a phenomenon referred to as the exponential-distance rule, EDR, characterized by the exponential decay rate, λ, see (Ercsey-Ravasz et al., 2013). The EDR held for the subset of areas investigated here (Fig. S5A), with λ=0.202/mm for distance through white matter, consistent with previous reports (Ercsey-Ravasz et al., 2013). Importantly, the EDR also held for the present FC data (Fig. S5A, all bands averaged for simplicity), yet with exponential decay rates that were substantially lower (0.01-0.08/mm; values per band and FC type reported in Table S2) (Fischer et al., 2018; Leopold et al., 2003; Nelson and Pouget, 2012).

Hence, a joint distance dependence might explain the observed correlation between FLNe and FC. Note that this would not explain the observed frequency dependence of the FLNe-FC correlation. Nevertheless, we investigated to which degree the FLNe-FC relation might be explained by distance. To this end, we performed a multiple linear regression (MR) with the dependent variable being FLNe, and the independent variables being the FC values for theta, beta, high-beta and gamma, and additionally the distance (as distance metric, we use distance on the cortical surface (Fig. 6) or distance through the white matter (Fig. S6), giving similar results and reported in Table S1). Note that this partial correlation also informs us about whether FLNe can be partly predicted by FC metrics. Figure S6E shows that FC alone (without distance information) is strongly predictive of FLNe, with explained variance (R^2^ full-model) ranging from 0.48 for GC to 0.56 for coherence (Table S1). This is interesting, because FLNe can currently not be obtained for the human brain, as it requires active retrograde transport and therefore the injection of retrograde tracer into the living brain. By contrast, FC and in particular GC can be obtained for the human brain, and GC has already been shown to relate to the SLN metric of anatomical connectivity patterns (Michalareas et al., 2016). The MR analysis revealed that all FC metrics were significantly predictive of FLNe for some of the frequency bands, and importantly, that this was the case when distance was included as an independent variable (Fig. 6A). Specifically, power correlation was significantly FLNe-predictive in the beta and gamma bands. Coherence and GC were FLNe-predictive in the gamma band. Note that the predictions of FLNe by those FC metrics was sufficiently accurate, such that when they were taken into account, the previously reported relation between distance and FLNe was no longer significant. Fig. 6A shows results obtained for distance measured on the cortical surface. When distance was measured through white matter, this explained slightly more FLNe variance, but overall, the pattern of results was highly similar (Fig. S6).

**Figure 6.**
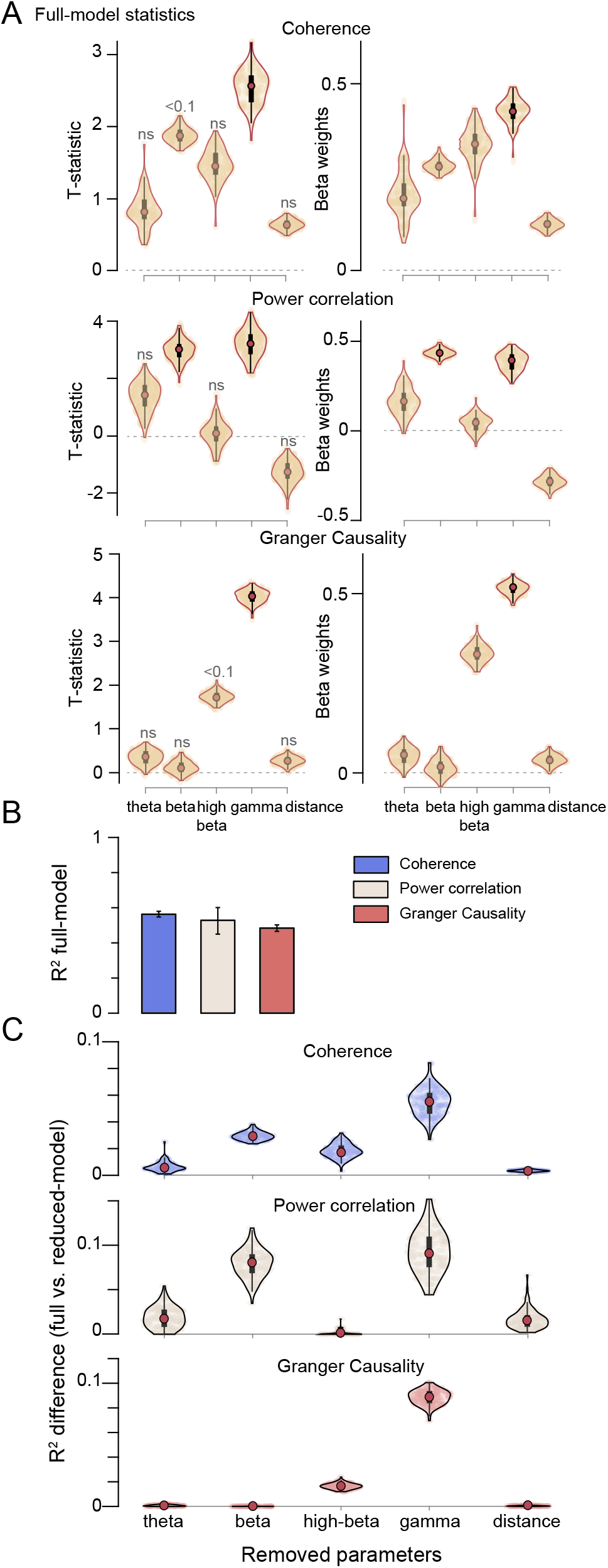
Multiple regression discloses distance as poor predictor of the structure-function relationship. (A) Violin plots of model estimates (t-statistic and beta coefficients, left and right column respectively) for each of the five variables considered i.e. frequency-bands FC and distance (from left to right) and for each of the three FC types (from top to bottom). (B) Total explained variance (R^2^) for the three models (color-legend, top-right). Mean ±99.9% confidence intervals from bootstrap estimates over trials. (C) Difference in total explained variance of the three models (same color code as in B) between the full- and the reduced-model, after removing the parameter listed on the x-axis. This estimates the contribution of each of the five parameters to the total explained variance.

To further investigate the differential FLNe-predictive power of the FC metrics in the different bands and distances, we performed the following analysis. We first determined R^2^ values for the full MR models, separately for power correlation, coherence and GC (Fig. 6B). We repeated this analysis after excluding either one of the frequency bands or distance as independent variable, i.e. we calculated R^2^-values for reduced models. Figure 6C shows the R^2^ difference between the full and the reduced model (similar to a stepwise linear regression approach); the x-axis lists the independent variable that had been removed, such that the corresponding y-axis values reflect the improvement in R^2^ value, when this variable is included. For coherence, all bands improved FLNe prediction, and the effect of distance was significant but very small. For power-correlation, the theta, beta and gamma bands improved FLNe prediction; with all bands included, distance still improved FLNe prediction. For GC, high-beta and particularly gamma improved FLNe prediction, and distance had essentially no effect. Importantly, for all FC metrics, removal of distance reduced R^2^ values by only relatively small amounts, less than the removal of most of the individual band-wise FC metrics. As above, distance through white matter had a larger effect, but overall, the pattern of results was highly similar (Fig. S6C). Also, complete removal of distance as independent variable left the overall pattern of results qualitatively unchanged; as expected, regression coefficients increased (Fig. S6D-F).

Note that these analyses revealed that all (log-transformed) FC metrics were linearly related to distance, leading to a multi-collinearity among the independent variables. We performed several analyses to control for this (Fig. S5B-C, Fig. S7, Table S1 and see SI for details).

### FLNe-FC relations depend on corresponding SLN values

The analyses so far suggest that FLNe partly determines FC values, with a specific spectral pattern. We had previously found that one aspect of FC, namely GC between two areas, is related to another aspect of AC, namely the feedforward/feedback characteristics of the corresponding connections captured by the SLN metric. When retrograde tracer is injected in area A, and the labeled cells are counted in area B, separately for the supragranular (Nsupra) and infragranular (Ninfra) compartments of B, then the SLN of the A-to-B projection is Nsupra / (Nsupra + Ninfra).

The larger the SLN metric, the more the corresponding projection is of feedforward type. Projections with SLN>0.5 can be considered feedforward, and projections with SLN<0.5 feedback. We previously found that if the SLN indicates that area B is higher in the hierarchy than area A, then theta- and gamma-band GC is stronger in the A-to-B (feedforward) than B-to-A (feedback) direction, whereas beta-band GC is stronger in the B-to-A (feedback) than A-to-B (feedforward) direction. Now, we will investigate, whether this SLN-GC relationship also affects the above described dependence of GC on FLNe.

To investigate this, we first selected two groups of projections, namely strongly feedforward projections, with SLN>0.7, and strongly feedback projections, with SLN<0.3. Within those two groups, we calculated the FLNe-related GC change for all frequencies (i.e. as in Fig. 5C, but split for SLN). For feedforward projections, the change spectrum showed peaks at theta and gamma, separated by a relative trough around beta (Fig. 7A, green line). For feedback projections, the change spectrum showed the strongest peak at high-beta, and a smaller one at gamma (Fig. 7A, black line). Figure 7B shows the corresponding scatter plots at the peak frequencies of each rhythm. For gamma GC, FLNe explained 48% (R^2^-value) of the variance in the feedforward, but only 37% in the feedback direction, whereas for beta GC, FLNe explained 15% in the feedback direction, but no significant part of variance in the feedforward direction (0.0001%, n.s.). The absence of a significant relation between FLNe and beta GC in the feedforward direction is also reflected in the beta-band trough (Fig. 7A, in green). We next determined the asymmetry index of the FLNe-related changes by taking the difference of the feedforward-minus the feedback-related spectrum and dividing by their sum (Fig. 7A, inset). This asymmetry index showed particularly pronounced negative values for beta and positive values for gamma, with much smaller effects for theta and high-beta. In order to test whether this result depended on the particular SLN cutoff (0.7/0.3), we repeated the same analysis for various cutoffs, and found that the observed effects generally showed a gradual dependence on SLN values (Fig. 7C).

**Figure 7.**
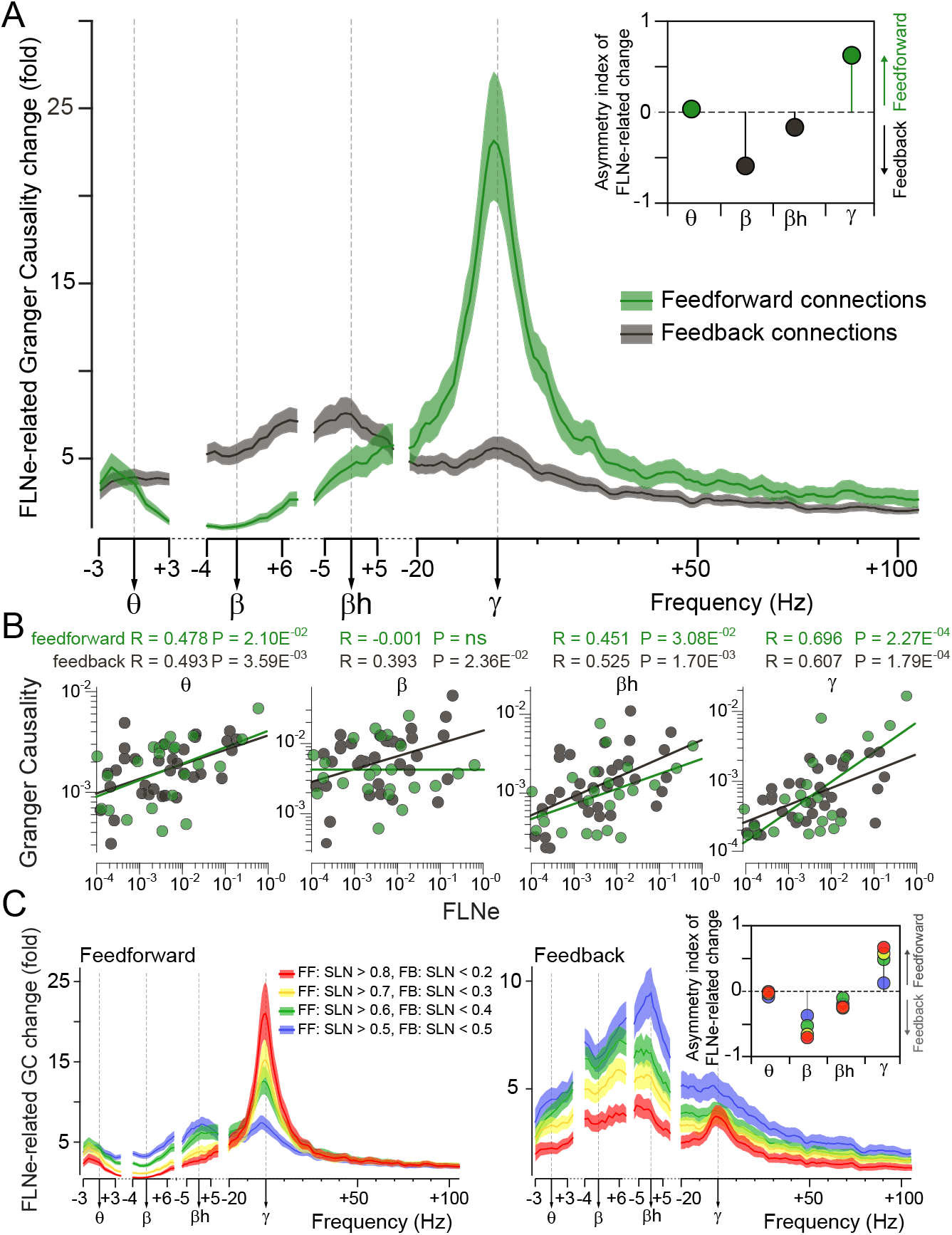
Anatomical influence on FC Strength depends on frequency and direction. (A) Frequency-resolved spectra of the FLNe-related change in GC (same as in Fig. 3), plotted separately for feedforward (in green, SLN ? 0.7) and feedback connections (in black, SLN ≤ 0.3), extracted from linear regression shown in (B). Inset (top right) displays asymmetry index (see Results) for all frequencybands. A positive (negative) asymmetry index indicates larger effect size in feedforward (feedback) direction. (C) Same as A, but varying the selection threshold for feedforward (left) or feedback (right) connections. Selecting more strongly feedback projections with lower SLN thresholds, resulted in lower FLNe-related GC changes (right panel). By contrast, selecting more strongly feedforward projections with higher SLN thresholds, resulted in lower FLNe-related GC changes for all frequency bands, except the gamma band, where this effect reversed. Note different axes scales for feedforward and feedback. Color code for varying SLN thresholds detailed in legend. Inset (top right) displays asymmetry index (see Results) for varying threshold (color code as in legend). Ordinate axes in (A) and (B) start from 1. All plots show mean over all trials ±99.9% confidence intervals from bootstrap estimates over trials.

### Mapping frequency-specific FC networks onto the anatomical core-periphery structure

We established that FC is related to both the strength (FLNe) and the feedforward character (SLN) of anatomical projections. The analysis of anatomical projections has integrated those two metrics and thereby demonstrated that areas can be arranged in a bow-tie structure: some areas are in the knot (the core) and others in the two fans (peripheries) of the bow-tie (Markov et al., 2013). Areas inside the core are densely interconnected and with strong (high FLNe) connections, whereas areas in the fans are connected less densely with areas in the core and with weaker connections to those areas. We found FC strength in the gamma frequency-band to dominate in the left fan areas of the bow-tie structure (Fig. 8A, S8A), i.e. areas sending predominantly feedforward projections to the core. FC strength at other frequencies was more distributed among core and periphery (Fig. 8B-D and S8B-D). Overall, FC strength was the strongest in the high-beta frequency-band for the core and in the beta frequency-band for right-fan areas of the bow-tie structure (Fig. 8C and S8C) i.e. areas sending predominantly feedback projections to the core.

**Figure 8.**
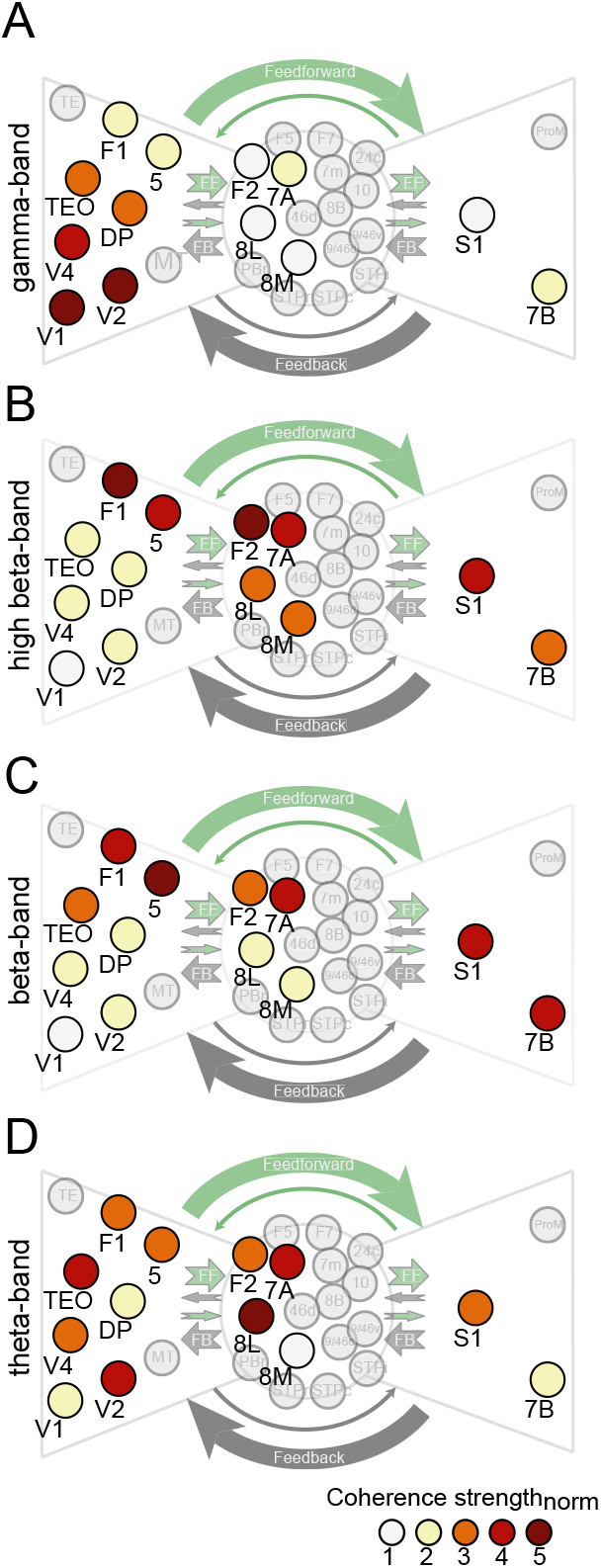
FC strength displayed on the AC-derived core-periphery structure. For cortical areas both recorded by the ECoG and used to build the core-periphery structure (highlighted areas) (Markov et al., 2013), the color code displays the respective area’s Coherence strength for the respective frequency band. Coherence Strength of an area was the average coherence of all sites of that area with all other sites in the ECoG (see Methods); it was averaged over monkeys before normalization into the range 1-5 (color scale, bottom right), separately for each frequency-band (A-D, gamma to theta). Same analysis for power correlation and GC shown in Fig. S8.

## Discussion

We used simultaneous ECoG-LFP recordings from 15 macaque cortical areas to compute FC matrices for 105 pairs of areas and the resulting strength topographies. This revealed pronounced differences across the four rhythms, yet also some commonalities. We then integrated this unique set of FC data with another unique set of AC data, namely the quantitative FLNe metric of the underlying anatomical projections between 14 of those areas, i.e. for 91 of those pairs of areas. Overall, we found that AC strength is a strong predictor of FC. Intriguingly, the influence of AC strength was weakest for power correlation, medium strength for coherence and strongest for GC. Furthermore, AC influenced coherence and GC in a frequency-dependent manner, with relative peaks for high-beta (coherence) and gamma (GC), and a relative trough for beta (both coherence and GC). Note that this trough is relative to neighboring frequencies, with the AC-FC correlation still significantly positive for beta. In addition, we show that FC strength is, in turn, a robust predictor of AC. GC and coherence at gamma, together with power correlation at gamma and beta, significantly predict the distribution of AC strength independently of the distance separating cortical areas. In addition, AC prediction by gamma GC and coherence is boosted by FC at beta (even though beta itself poorly predicts FLNe), suggesting that part of the explained FLNe variance (R^2^ full-model) is due to cross-frequency interaction between these bands (Richter et al., 2017). We then investigated the dependence of GC on AC separately for anatomical feedforward and feedback projections. The strength of feedforward projections was primarily predictive of GC at gamma. Conversely, GC at beta was only significantly predicted by the strength of feedback but not feedforward projections. This might be related to our previous finding among visual areas, that GC at gamma is stronger in the anatomically defined feedforward direction, and GC at alpha/beta is stronger in the feedback direction (Bastos et al., 2015; Michalareas et al., 2016). To summarize, we show that AC determines up to half of the variance in FC, and conversely that more than half of the variance in AC can be predicted by FC. These values are likely underestimated, because we used FC and AC values from separate sets of animals, and AC strengths vary across individuals by about an order of magnitude (Markov et al., 2014; Markov et al., 2011). Even among AC values, there was additional inter-individual variability, because data for different projections originated partly from different animals. Furthermore, FC values are known to be strongly modulated by stimuli, state and task, and including FC values from all those conditions should further improve the ability to predict AC values.

One potential limitation of the present study is that any spectral decomposition of the LFP is imperfect, such that energy can partly leak across neighboring bands. In particular, the beta band typically has relatively strong power for many of the recorded brain areas, and it could partly leak into the theta and high-beta bands. Such leakage would pull the results for neighboring frequency bands towards those for beta, which might partly explain some of the overlap in strength topographies. However, among the four investigated rhythms, coherence and GC at beta were least affected by AC strength meaning that the relatively stronger influences of AC strength on the neighboring theta and high-beta bands might have been under-estimated, but they are most likely not created by spectral leakage.

Several previous studies have investigated large-scale patterns of rhythmic synchronization, primarily with non-invasive methods in human subjects. MEG and source-level FC metrics have been used to investigate several cognitive functions and led to the association of specific high-order functions to specific brain-wide networks synchronized at different frequency-bands (Gross et al., 2004; Gross et al., 2002; Hipp et al., 2011; Kujala et al., 2007; Rouhinen et al., 2020; Siegel et al., 2008). Besides these task-related networks, also intrinsic networks of the resting brain have been investigated with source-projected MEG and revealed well-characterized resting-state networks through power correlation at different frequency-bands (de Pasquale et al., 2010; de Pasquale et al., 2012). However, the interpretation of FC metrics based on non-invasively recorded signals is complicated by the fact that those signals always reflect mixtures of many brain sources, and source-separation methods cannot fully undo this mixing. This has e.g. been demonstrated for the beta-band synchronization between different components of the motor system: While those components can be separated in an analysis of coherence of brain source to the separately recorded electromyogram, they cannot be distinguished in an analysis of coherence among brain sources (Schoffelen et al., 2008). However, a few studies have at least mitigated the problem of signal mixing. The use of Granger causality on source-projected MEG data seems to reduce the problem of signal mixing, probably because Granger causality reflects causal, i.e. time-delayed, interactions, and explicitly discards instantaneous interactions resulting from signal mixing (Michalareas et al., 2016). In addition, the present results suggest that GC is a valid measure of interareal synaptic influence as it captures much of underlying connection strengths. In the analysis of power correlation, signal mixing can be effectively reduced by prior orthogonalization of the band-passed signals (Hipp et al., 2012). This approach dramatically reduces the power-correlation values observed between separate brain sources, and reveals bilaterally symmetric interhemispheric power correlation reminiscent of fMRI-based FC studies. When combined with fMRI BOLD signal correlation analysis, the relation between electrophysiological and hemodynamic correlation structures extended over a broad frequency range from 2 to 128 Hz, with substantial variability across cortical space in the frequency band with the tightest link to BOLD correlation (Hipp and Siegel, 2015). Thus, this approach provides intriguing links between two non-invasively measured metrics of connectivity. However, the interpretation of fMRI BOLD signal correlations is difficult due to the indirect and temporally sluggish link between the hemodynamic BOLD signal and the underlying neuronal activity (Logothetis et al., 2001). A very different approach to using MRI for connectivity analysis capitalizes on diffusion-tensor imaging (DTI), which is based on the anisotropic diffusion of water in highly organized fiber bundles of the brain (Basser et al., 1994). However, DTI is limited in assessing anatomical connectivity, and the correlation between log-transformed DTI-based tractography and retrograde-tracer based connection weights is merely 0.59, and drops to 0.22, when joint distance dependence is taken into account (Donahue et al., 2016).

The limitations of non-invasive approaches can be overcome primarily in animals, through invasive electrophysiology and tracer-based quantification of projection strengths. Similar to MEG-based studies of synchronization, most animal studies of interareal synchronization were so far focused on modulations by sensory stimulation, attention, or other cognitive factors (Singer, 2018). In particular, spatially selective visual attention strongly modulates gamma-band synchronization between prefrontal and occipital cortices (Gregoriou et al., 2009), and also between lower and higher visual areas (Bosman et al., 2012; Grothe et al., 2012; Richter et al., 2017; Rohenkohl et al., 2018). Interareal influences at gamma versus alpha/beta subserve distinct functions in bottom-up and top-down communication: interareal synchrony is stronger in lower frequencies during top-down attention and in higher frequencies during bottom-up attention (Buschman and Miller, 2007). In hierarchical prediction paradigms, gamma and alpha/beta bands convey prediction errors and prediction updates, respectively (Chao et al., 2018). Likewise, predictable stimuli induce power and FC primarily in alpha and beta band and related to deep layers, whereas unpredictable stimuli induce the same in gamma and related to superficial layers (Bastos et al., 2020). However, the relation between rhythmic synchronization and the underlying AC has been studied within few brain regions. In cat visual cortex, gamma-band coherence is particularly weak between monocular neurons in strabismic animals (König et al., 1993), which largely lack lateral connections (Löwel and Singer, 1992). Similarly tuned regions of cat area 17 are linked by the supragranular daisy network (Martin et al., 2017) and display enhanced coupling (Cossell et al., 2015; Ts’o et al., 1986). In squirrel monkeys, FC patterns of synchronized firing within and between somatosensory areas are consistent with their AC and rs-fMRI patterns (Wang et al., 2013).

The analysis of anatomical connectivity, derived from tracer injections in animals or from MRI BOLD signal correlations or DTI, has provided crucial insights into the large-scale connectome of the primate brain. The present work reveals the differential relation between anatomical connectivity and rhythm-based functional connectivity in different frequency bands, and it presents band-wise connectomes for large parts of the brain surface. Intriguingly, recent evidence suggests differential links between the band-wise human connectomes and individual behavioral measures (Becker and Hervais-Adelman, 2020).

There are several indications that structural and functional connection weights impact across multiple scales. At the level of the single-neuron, investigations report a skeleton of strong connections in a sea of weak connections and weights follow a lognormal distribution (Buzsáki and Mizuseki, 2014; Song et al., 2005). The strength of synaptic connections appears to determine neuronal firing since the few strong connections occur between neurons with similar receptive field properties and with the most correlated response, while weak connections are found between uncorrelated neurons with dissimilar receptive fields (Cossell et al., 2015). At the mesoscale, interareal connection weighs show a lognormal distribution (Gămănuţ et al., 2018; Markov et al., 2011; Oh et al., 2014; Wang et al., 2012). The functional significance of the numerous weak connections remains essentially a mystery both for single-neuron and interareal connectivity. The present results indicate that weak long-distance connections carry longdistance effects through synchronization in distinct frequency bands.

## Methods

### Neurophysiological Recording Techniques and Signal Preprocessing

All procedures were approved by the animal ethics committee of Radboud University (Nijmegen, The Netherlands). Neuronal signals were recorded from the left hemisphere in two male rhesus monkeys using subdural ECoG grids consisting of 252 electrodes (1 mm diameter), which were spaced 2-3 mm apart. All analyses use local bipolar derivations, i.e. sample-by-sample differences, between neighboring electrodes (for details, see (Bastos et al., 2015)). We refer to bipolar derivations as “(recording) sites”. Here, we selected areas and the corresponding site pairs, if they were also injected with retrograde tracers. This resulted in a total of 14 areas, which were electrophysiologically recorded in two macaques, and injected with tracers in a separate cohort of 26 macaques. The list of selected areas is: V1, V2, 8L, V4, TEO, DP, 8M, 7A, S1, 5, 7B, F1, F4, F2. To minimize volume conduction effects, we excluded site pairs with an inter-site distance (along the dural surface) of less than 4 mm from the calculation of interareal averages. Note that this corresponds to the diameter of an anatomical macrocolumn: Anatomical tract-tracing studies have shown that 95% of intrinsic connections are located within a distance of 1.9 mm centered on the injection site (Markov et al., 2011). Site-pairs containing artefacts were rejected after visual inspection. Signal amplification, filtering, digitization and preprocessing are described elsewhere (Bastos et al., 2015). In short, only trials with a correct behavioral report were used, pooling both attention conditions (9 sessions in Monkey1 and 14 sessions in Monkey2). Power line artifacts at 50 Hz and its harmonics up to the Nyquist, as well as screen refresh-rate artefacts (120Hz) were then estimated and subtracted from the data using a Discrete Fourier Transform. The analysis described here used the period in the trial, during which stimuli were presented, and the monkey paid attention to one of them, called the post-cue period (for details, see (Bastos et al., 2015)).

### Visual Stimulation

Stimuli and behavior were controlled by the software CORTEX as described previously (Bastos et al., 2015). In short, during the visual attention task, monkeys fixated a central spot and released a bar when the behaviorally relevant stimulus underwent a shape change i.e. one the two gratings that color-matched the cue (central spot) would have its stripes undergoing a gentle bend lasting 0.15s.

### Data Analysis General

All analyses were performed in MATLAB (MathWorks) using FieldTrip (www.fieldtriptoolbox.org) (Oostenveld et al., 2011) and custom scripts. Data epochs were multitapered using three Slepian tapers and Fourier-transformed (Mitra and Pesaran, 1999). For epoch lengths of 1 s, the spectral resolution was 1 Hz, and the multitapering resulted in a spectral smoothing of ±1.5 Hz. The Fourier transforms were the basis for calculating the FC spectra, i.e. coherence spectra, spectra of power correlation across trials (Lewis et al., 2016) and the GC spectra. GC spectra were estimated through non-parametric spectral matrix factorization of the cross-spectral density matrices (Dhamala et al., 2008).

### Retrograde tracing database

Acquisition and analysis of the anatomical data set has been reported in (Markov et al., 2014a). Updates, atlases and additional information concerning the anatomical data set that was used for this work is available at www.core-nets.org. We used the fraction of labeled neurons (FLNe, defined in the Resultssection) to quantify AC strength. We used the proportion of supragranular labeled neurons (SLN, also defined in the Resultssection) to quantify the feedforward or feedback nature of an anatomical projection. Furthermore, we used interareal white-matter distances. These metrics were obtained for all of the 91 pairs of areas, which were also covered by the ECoG grids i.e. between the 14 cortical areas V1, V2, 8L, V4, TEO, DP, 8M, 7A, S1,7B, 5, F1, F4 and F2.

### Registration of ECoG grids to individual and template volumes

The anatomical MRI of each subject was spatially coregistered with the 3D positions of electrode locations using the FieldTrip toolbox (Oostenveld et al., 2011). These 3D positions were obtained by projecting the 2D positions (from high-resolution intraoperative photographs, using the sulci for alignment, see (Bastos et al., 2015)) onto each individual brain surface using the iso2mesh toolbox (http://iso2mesh.sourceforge.net). Each individual anatomical MRI was coregistered to the F99 template brain containing anatomical atlases information (Van Essen, 2012), using linear and non-linear coregistration with FSL (Smith et al., 2004) (Fig. S1) and further aligned and wrapped to the INIA19 macaque brain surface (Rohlfing et al., 2012). The transformation matrix was then applied to the volume of each ECoG electrode positions in order to i) assign each site to the underlying cortical area similarly as done in (Bastos et al., 2015) and ii) combine the two ECoG grids on this template surface (Fig. 1B) in order to create FC strength maps (Fig. 2–3), after averaging overlapping parts of the two ECoG grids. We calculated distances separating recording sites along the dural surface using the fast-marching toolbox in MATLAB (MathWorks).

### Statistical testing

Wherever possible, data from both monkeys were combined. The combined results amount to a fixed-effect analysis, allowing an inference on our sample of two animals. A frequency-dependent smoothing with a span of ±1% of the respective center frequency was applied to frequency spectra for data visualization. Results presented are averages over interareal pairs and over all epochs. 99.9% confidence intervals were estimated from a bootstrap procedure over epochs as described in (Efron and Tibshirani, 1994). One-hundred bootstrap samples were calculated for each area pair and each monkey, before averaging over monkeys. Averaging over monkeys was done after peak-alignment for each of the four frequency bands of interest. The correlation with FLNe was then performed and relevant statistics extracted, i.e. rho, p-value and slope. We additionally extracted the FLNe-related FC change as the difference between predicted FC values for minimal and maximal FLNe by the linear regression. Values have been logged (base10) before performing linear regression, essentially to account for the widespread variance in FLNe and FC values (several orders of magnitude as depicted in Fig. 5B). Therefore, the FLNe-related FC change directly reflects a fold change. To further capture the complexity of the structure-function relationship, we performed multiple linear regression (MR) of FLNe by FC for all frequency bands of interest (Fig. 6). Because FLNe obeys an exponential distance rule (Ercsey-Ravasz et al., 2013), we verified that FC follows a similar exponential decrease with either interareal distance as measured between site-pairs along brain surface or through white-matter distance (Fig. S6). Decay rates were calculated using normal log as described in (Ercsey-Ravasz et al., 2013). By integrating distance in the regression model, we controlled for this potentially confounding variable and provide the partial correlation coefficients. However, in parallel to the expected bias reduction, the risk of data collinearity could in turn potentially reduce the precision of model estimates. We thus controlled for the non-violation of the ordinary least square assumption (Fig. S5B) and verified that variance inflation factors (VIFs) remained below critical levels, in particular for variables with significant model coefficients (Fig. S5C). The VIF for a given predictor variable indicates the degree to which collinearity potentially inflates the standard error of its coefficient estimates, thereby reducing statistical power and warranting caution in the interpretation for this predictor (VIFs start at 1 meaning no correlation between predictor variables and any other; values between 2.5 and 5 indicate moderate correlation but not warranting corrective measures; values above 5 indicate a critical level). Importantly, even in the presence of correlation between variables, multicollinearity does not compromise the interpretation of MR coefficients provided this is done on grounds of outcome from analyses allowing assessment and control for collinearity, e.g. considering dominance or relative importance of partial regression coefficients. Hence, in addition to structure coefficients – measured already independently of collinearity and dividing each variable’s contribution to the multiple regression effect – we measured direct effects of predictors and shared effects between predictors through ‘unique’ and ‘common’ coefficients calculated from commonality analysis (CA, performed with y-hat package under R, https://www.R-project.org/). For each predictor, the squared structure coefficients (rs^2^) characterize the shared amount of variance with – or the individual contribution to – the multiple regression effect (R^2^), and therefore should be interpreted as the amount of explained effect rather than explained variance of the dependent variable. In case of multicollinearity, CA provides the very useful direct and shared coefficients of total explained variance (R^2^) to each subset of predictors from all possible subsets regression. It additionally allows identification of so-called ‘suppressor’ variables, through negative common coefficients which estimate the amount of predictive power lost by other predictors when removing the considered variable(s) from the MR model. Direct or ‘Unique’ effects are comparable to change in the multiple coefficient of determination from squared semi-partial correlation after inclusion of a variable at last position of a hierarchical regression. Formulas for direct and shared components of a predictor variable Xi from a model with n predictors are, respectively Ui=-(1-Xi)XjXk… Xn) and Ci=-(1-Xi)(1-Xj)(1-Xk)…(1-Xn). Other relative importance measures considered and reported in Table S1 are Effect Size (for adjusted R^2^), General Dominance weights (GenDom – average of overall conditional dominance weights i.e. additional contribution to multiple R^2^ computed in all possible predictors combination comparisons) and Relative Importance Weights (RIW – proportional contribution to multiple R^2^ after correcting correlation amongst predictors). Dominance analysis ranks predictors based on explained variance for all pairwise comparisons and minimizes the contribution of predictors in presence of collinearity. Thus, conclusions from GenDom and RIW should be consistent. Importantly, the sum of all weights equal the multiple R^2^ of the MR model for both.

Finally, similar to the principle of stepwise MR, we calculated the individual variables contribution to the R^2^ of the full-model by comparing the latter to the R^2^ of the reduced model after removing this variable (Fig. 6C and S6C,F). We performed regression analyses for each of the 100 samples per predictor variable estimated from bootstrap over epochs, except for distance measures which do not change across trials.

All violin plots use bootstrap estimates over trials, their shape along the y-axis uses kernel density estimate with a self-optimized bandwidth of the density kernel (https://github.com/bastibe/Violinplot-Matlab/blob/master/violinplot.m).

Statistical significance for average FC strength maps was calculated by comparing actual values for each site to values obtained from 1000 permutations of labels across sites, using a controlled false discovery rate of 20% with an alpha of 0.05 (Korn et al., 2007). By doing so we compared the topography of frequency-specific maps to those obtained from a random graph with original FC values and the same number of nodes and edges.

## Author Contributions

C.A.B. and P.F. designed the experiments; C.A.B. trained the monkeys and recorded the electrophysiological data; J.V., M.V., A.M.B., C.L. and C.A.B. wrote analysis programs; J.V. performed analyses with the help of M.V. and with the advice of P.F.; H.K. provided the anatomical data; J.V., M.V., H.K. and P.F. wrote the paper in collaboration with A.M.B., C.L. and C.A.B.

## Acknowledgments

This work was supported by DFG (SPP 1665 FR2557/1-1, FOR 1847 FR2557/2-1, FR2557/5-1-CORNET, FR2557/6-1-NeuroTMR, FR2557/7-1-DualStreams to P.F.), EU (HEALTH-F2-2008-200728-BrainSynch, FP7-604102-HBP, FP7-600730-Magnetrodes to P.F.), a European Young Investigator Award to P.F., National Institutes of Health (1U54MH091657-WU-Minn-Consortium-HCP to P.F.), the LOEWE program (NeFF to P.F.).

The authors would like to thank Jarrod Dowdall for helpful support at some stages of the analysis and Georgios Spyropoulos for insightful discussions on earlier versions of the manuscript.

## Competing interests

P.F. and C.M.L. are beneficiaries of a license contract on thin-film electrodes with Blackrock Microsystems LLC (Salt Lake City, UT), and P.F. is member of the Scientific Technical Advisory Board of CorTec GmbH (Freiburg, Germany), and managing director of Brain Science GmbH (Frankfurt am Main, Germany).

## Materials & Correspondence

All data are presented in the main and supplementary figures and further available upon request. Correspondence should be addressed to.

## Supplemental Information

### Control analyses for the presence of collinearity in multiple linear regression

In order to investigate whether FLNe can be predicted by FC metrics independently of distance, we performed a partial correlation in the form of a multiple linear regression (MR). The dependent variable being FLNe, and the independent variables being the FC values for theta, beta, high-beta and gamma, and distance (as distance metric, we use distance on the cortical surface (Fig. 6) or distance through the white matter (Fig. S6), giving similar results and reported in Table S1).

These analyses revealed that all (log-transformed) FC metrics were linearly related to distance, leading to a multi-collinearity among the independent variables that may have affected the MR analysis. To investigate the severity of this, we performed several control analyses.

First, we plotted the residuals of the MR as a function of the predicted FLNe values separately for each FC metric (Fig. S5B). This revealed no systematic relationships, i.e. no indication of relevant unobserved (hidden) variables.

Second, we determined the variance-inflation factor (VIF), separately per FC metric and frequency band (Fig. S5C). These values were below the critical threshold (see Methods) for all combinations of FC metrics and frequency bands that had been found significantly predictive of FLNe in the previous analysis (Fig. S5C).

Third, we performed an analysis of structural coefficients and general dominance (Fig. S7, see Methods). We computed squared structure coefficients (rs^2^), general dominance (GenDom) and relative importance weights (RIW), as well as direct and shared effects for each variable, including distance (Table S1, Fig. S7). Results demonstrate that FC in the gamma band is the dominant variable, with the strongest direct effect and the highest relative importance in MR models predicting FLNe, with or without partial regression of distance. Commonality analysis identified a substantial proportion of the regression effect caused by suppression in MR models, consistently involving FC in the beta-frequency band (Fig. S7, sum of negative shared coefficients for Coherence, Power Correlation and GC: 9.73%-29.4%-15.03% without distance, 17.64%-29.75%-17.79% with surface-distance and 12.97%-27.74%-15.98% with WM-distance). The amount of overall suppression caused by beta-band FC in the model was strongest when combined with FC in the gamma frequency-band (insets Fig. S7A-C) suggesting subtle involvement of cross-frequency interactions in the observed regression effect. Furthermore, in all models, we observed that beta-band FC contributed the least to the MR effect (i.e. lowest rs^2^). Hence, even if poorly predictive of FLNe, the main contribution of beta-band FC to the MR effect was to increase FLNe-predictive power of the remaining variables and mostly for gamma-band FC.

**Figure S1.**
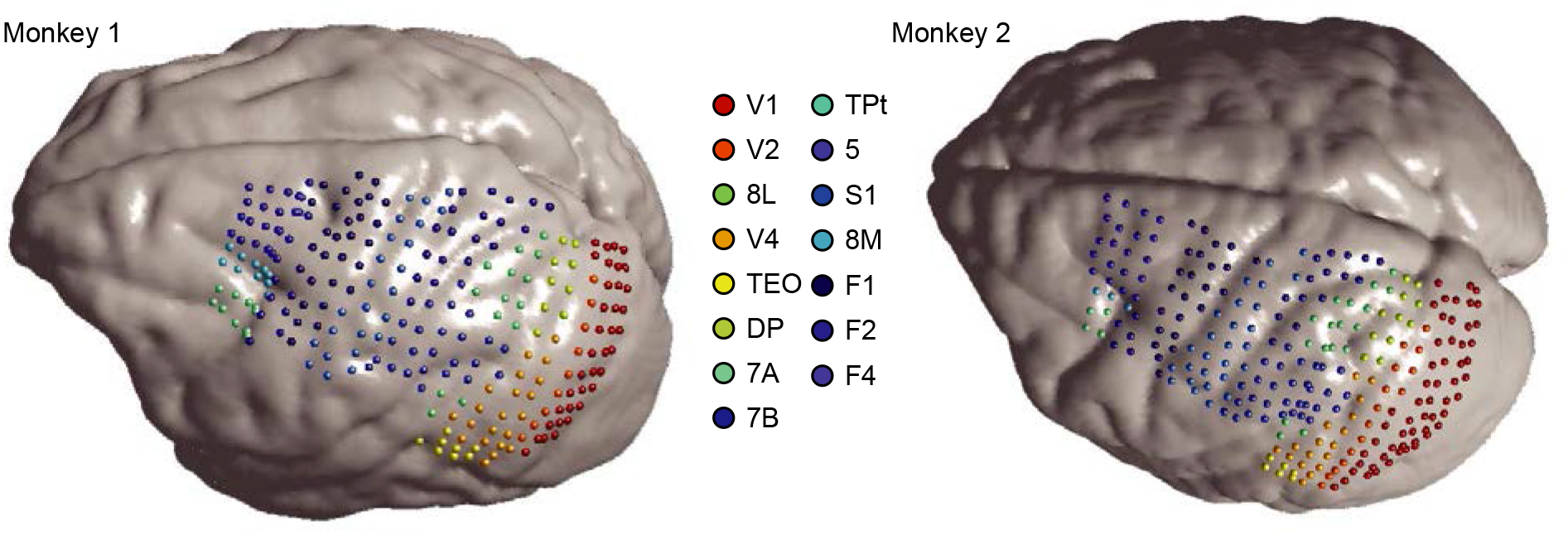
Individual recording sites. ECoG grids registered to brain surface reconstructions from individual MRIs. Sites are color coded according to underlying cortical areas, as indicated by the legend in the middle.

**Figure S2.**
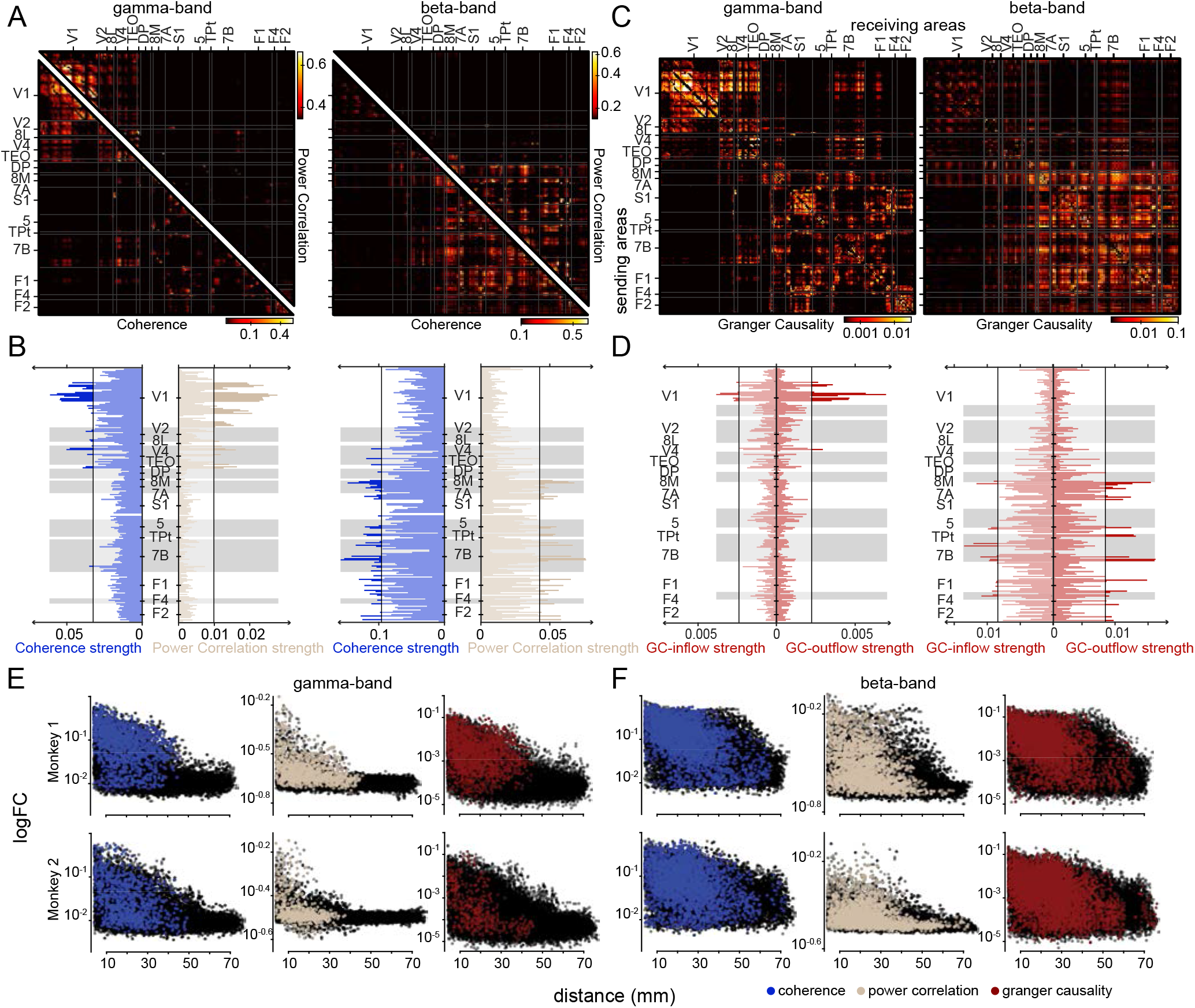
Functional connectivity matrices and area-wise strength in the gamma- and beta-band. **A-D.** Same conventions as in **Fig. 2** but for Monkey2. **E.** FC (log scale) relative to inter-site distance over the dural surface for Monkey 1 (upper row) and Monkey 2 (lower row) in the gamma-band. **F.** Same as in E but for FC in the beta-band. All possible site-pairs shown in black, and significant site-pairs colored according to the FC metric considered (legend, lower right). Significance determined by comparison to random graphs with original FC values and the same number of nodes and edges (see Methods).

**Figure S3.**
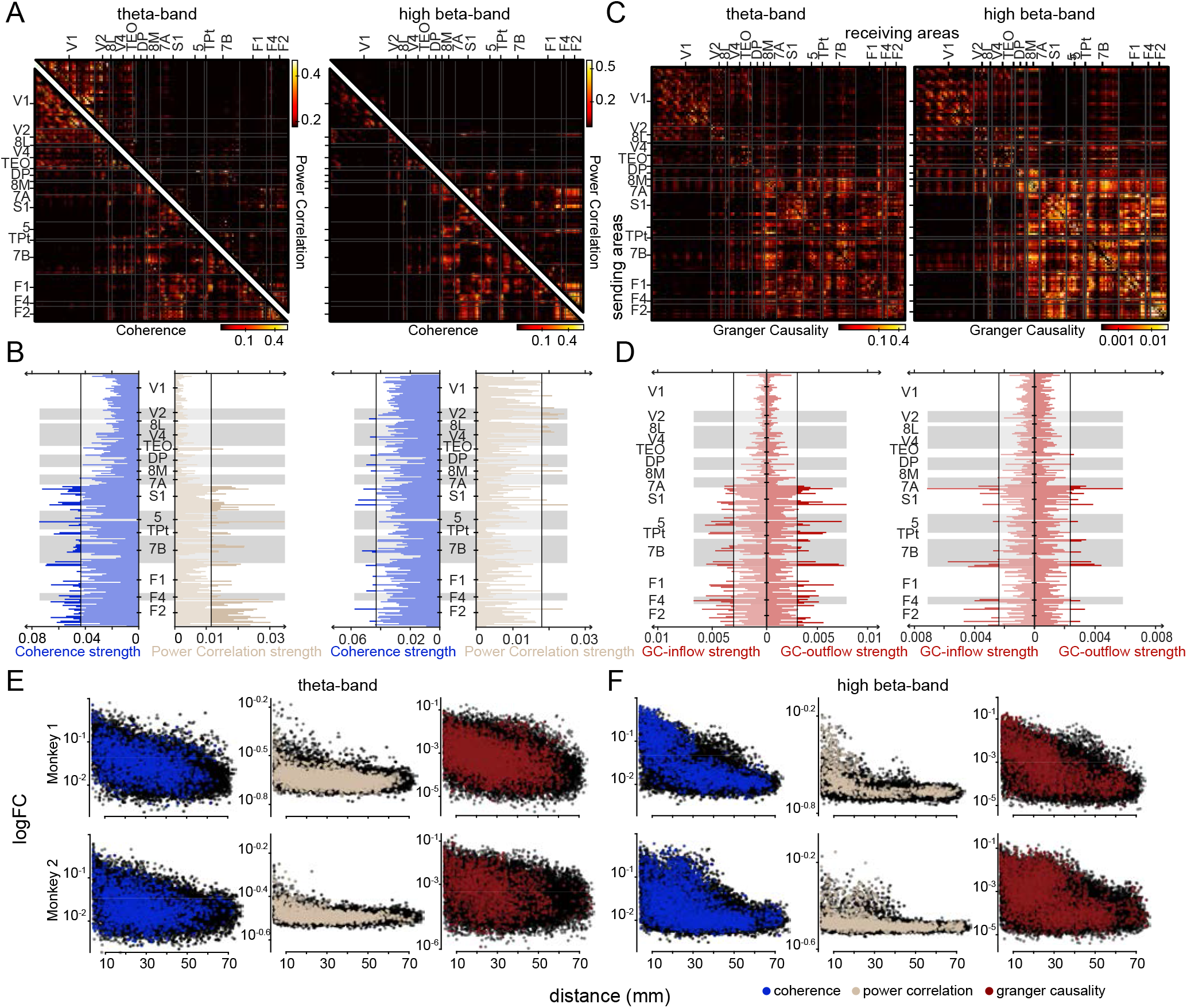
Functional connectivity matrices and area-wise strength in the theta- and high beta-band. **A-D.** Same conventions as in **Fig. 3** but for Monkey2.**E-F.** Same conventions as in **Fig. S2E-F** but for theta (**E**) and high-beta (**F**).

**Figure S4.**
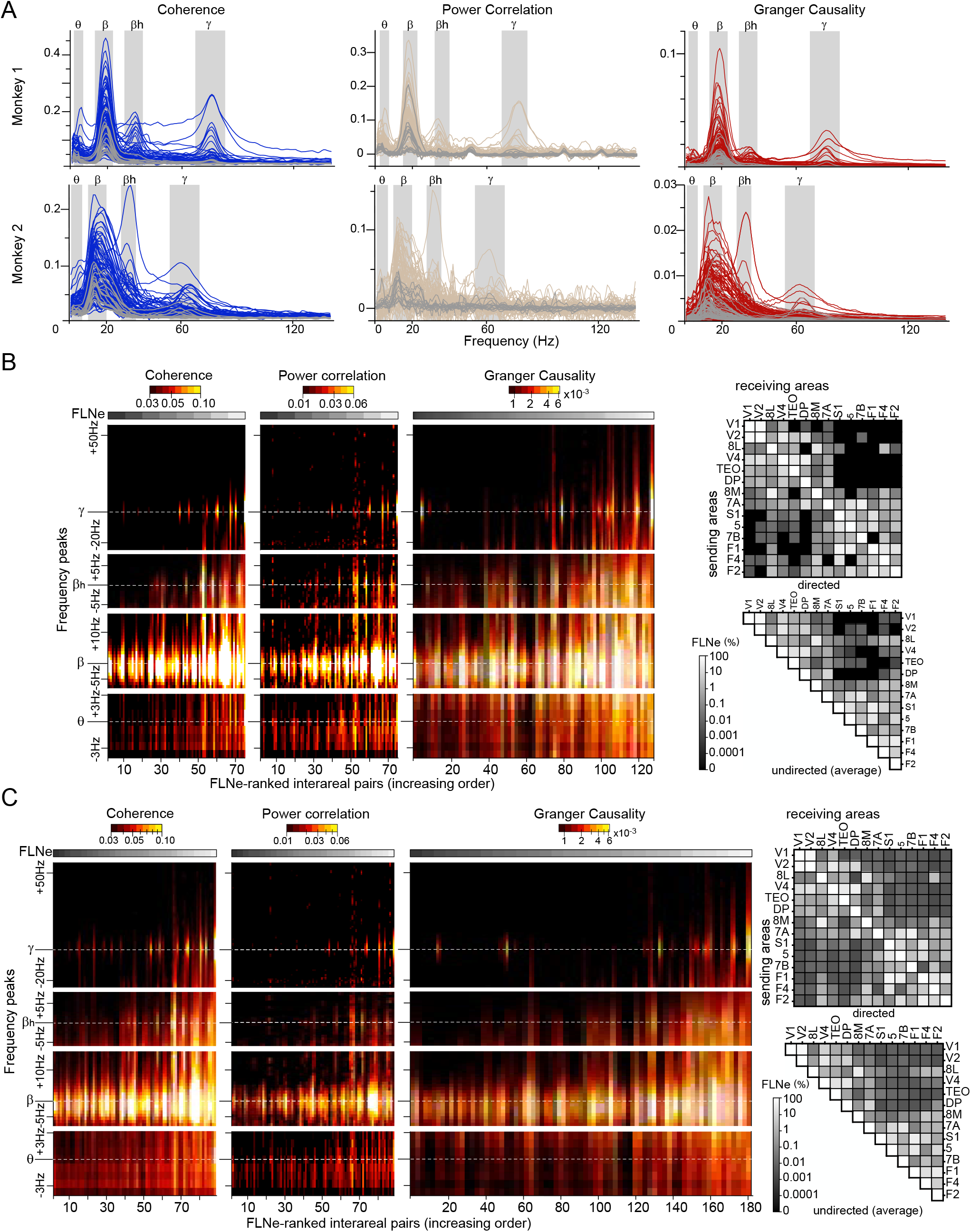
FC spectra and their relation to FLNe. **A.** Same conventions as in **Fig. 1C**. Interareal pairs for which no FLNe was reported i.e. no identified anatomical connection, are shown in light grey. **B.** Same as Figure 4D, but including FLNe values based on less than 10 neurons, while still excluding FLNe values of zero. The corresponding FLNe matrices are shown on the right. **C.** Same as Figure 4D, but including FLNe values based on less than 10 neurons, and replacing FLNe values of zero by estimates assuming Poisson distributions fitted to neuron counts from non-zero FLNe distributions. The corresponding FLNe matrices are shown on the right.

**Figure S5.**
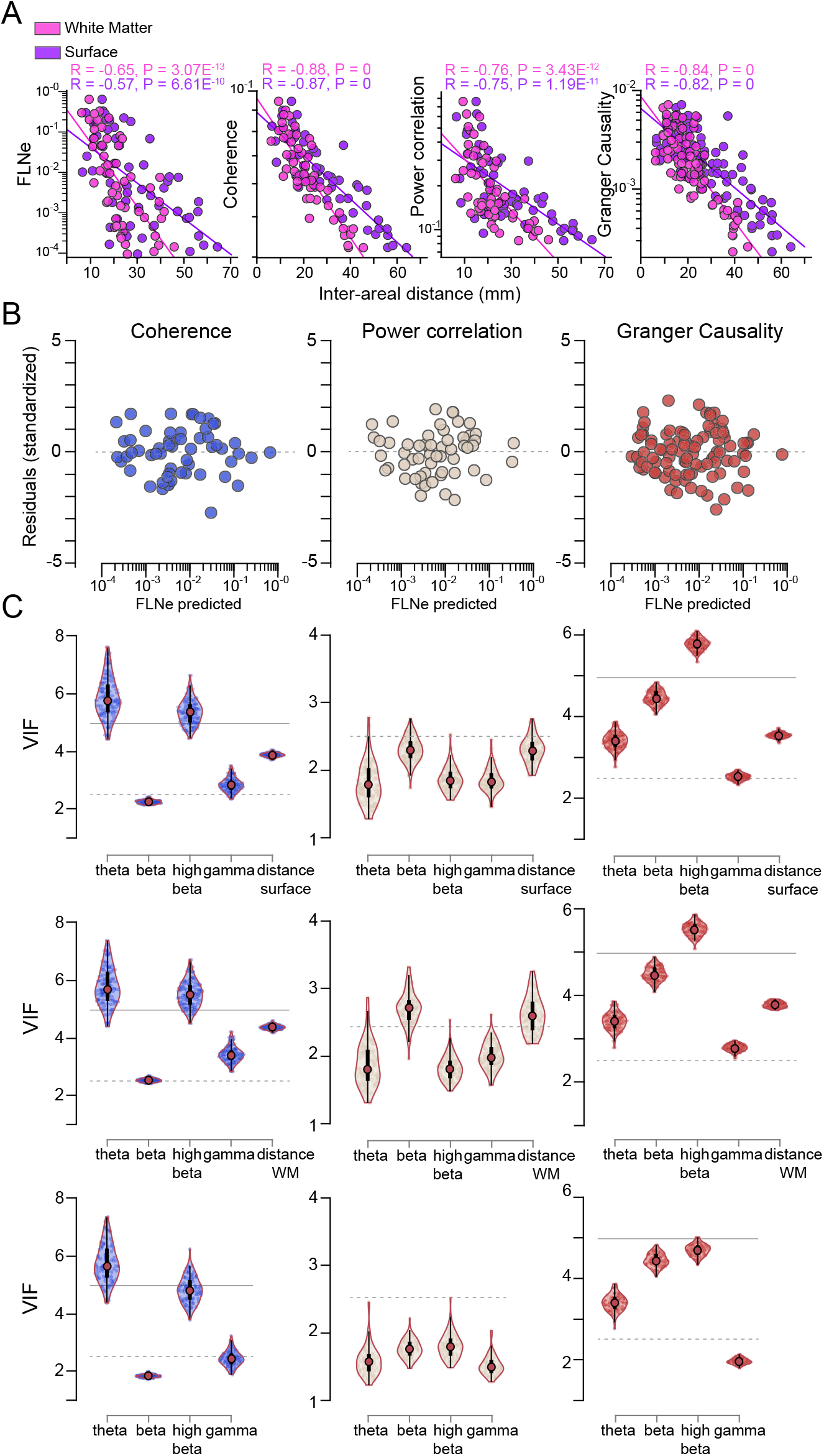
Anatomical and functional connectivity correlate with interareal distance. **A.** Linear regression of log(FLNe), log(Coherence), log(Power correlation) and log(Granger Causality) (left to right) against interareal distance on the brain surface (purple) or through white-matter (pink). FC values were averaged over the four frequency band of the two monkeys. Individual decay rates are presented in Table S2. **B.** Scatterplots of residuals against predicted variable verify heteroscedasticity for each FC metric. **C.** Variance Inflation Factor (VIF) calculated for each predicting variable of the three models, with interareal distance on the brain surface (top) or through white-matter (WM, middle) as an additional predicting variable or not (bottom). Dashed (VIF = 2.5) and plain lines (VIF = 5) show moderate correlation range still below critical level.

**Figure S6.**
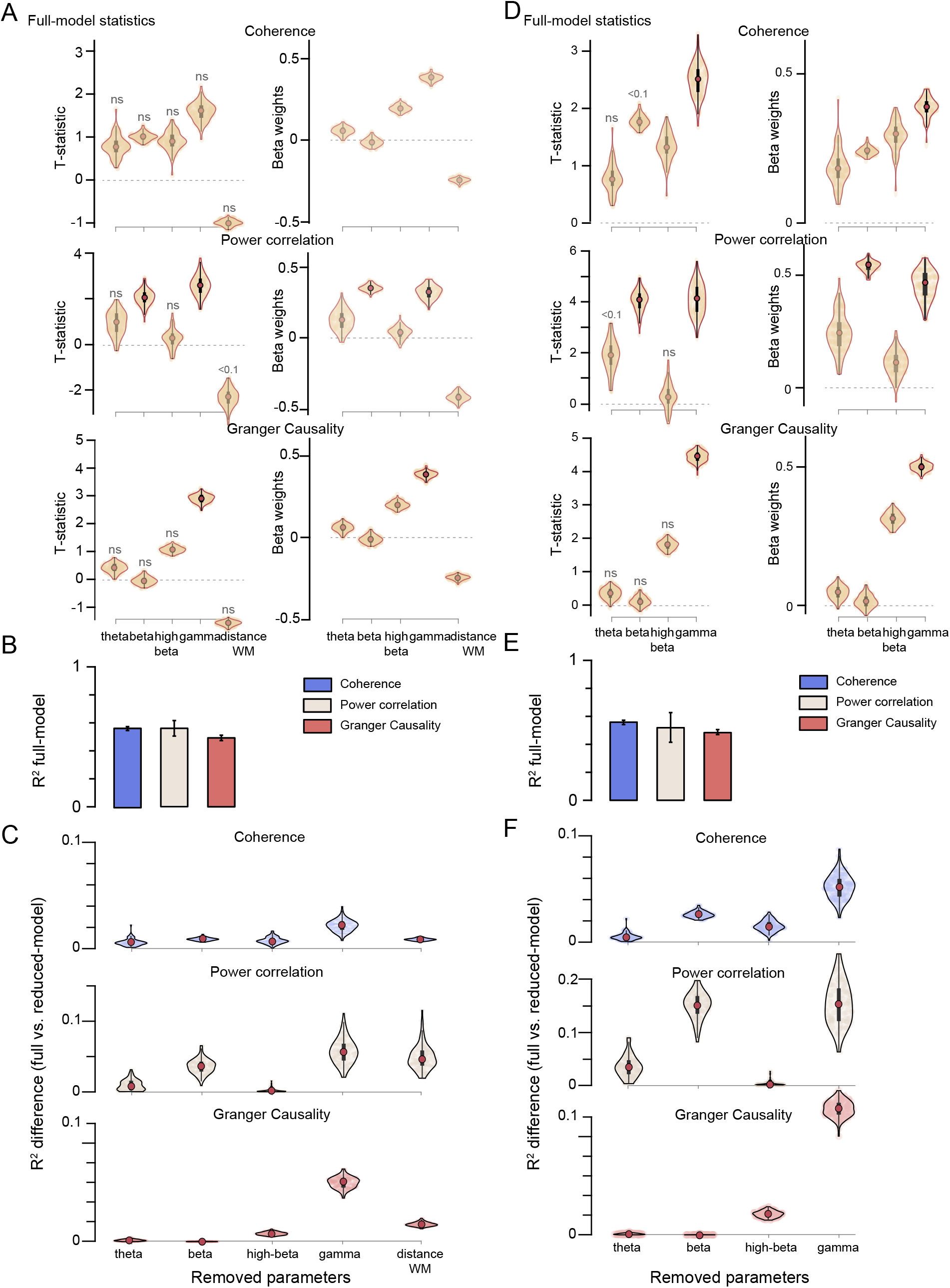
Multiple linear regression with or without interareal distance as additional predictor. **A-C.** Same as Figure 6A-C, but using interareal distance through white-matter (WM). **D-F.** Same as Fig. 6A-C, but not including interareal distance as predicting variable.

**Figure S7.**
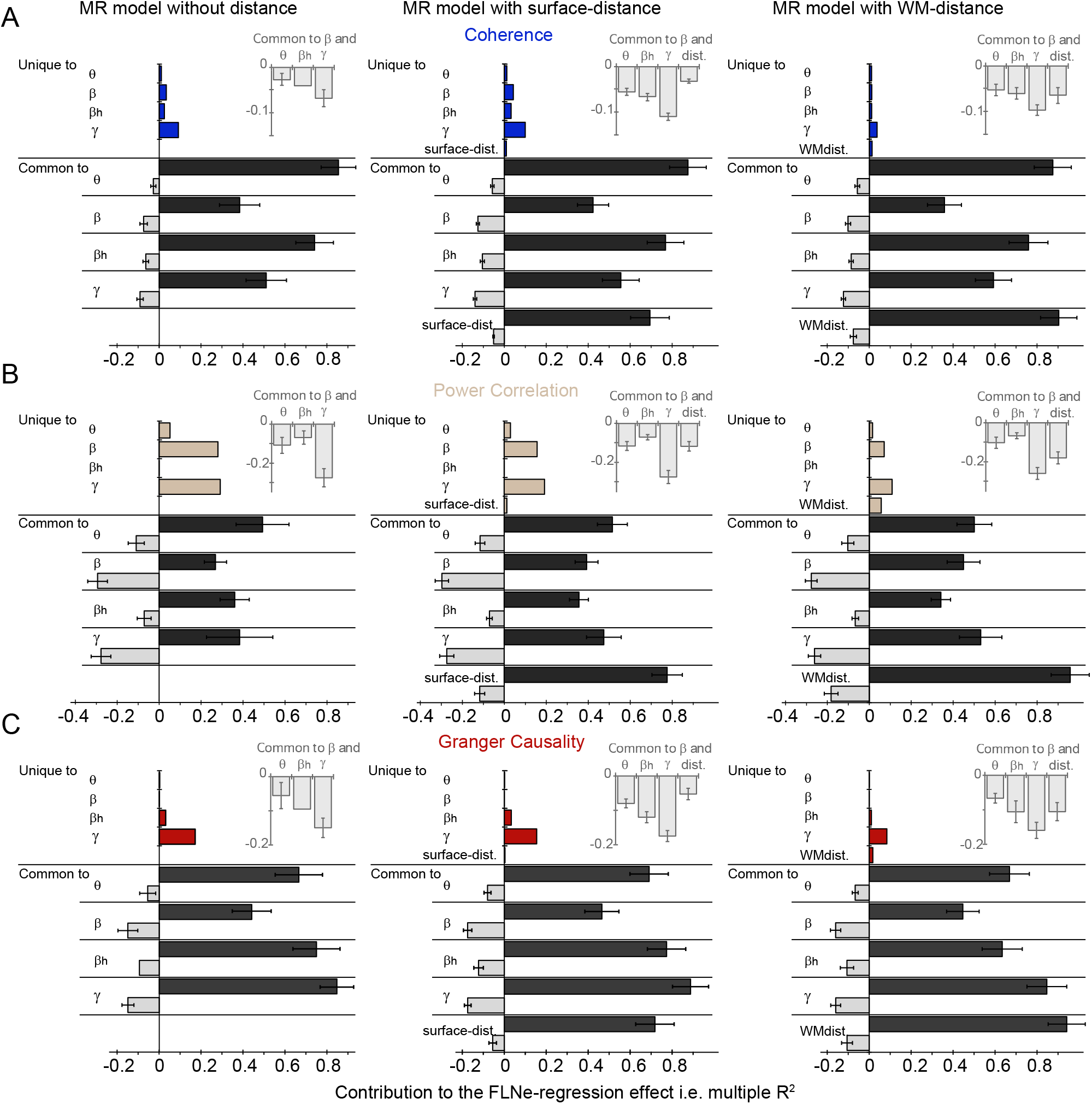
Commonality analysis summary. Unique (*Direct*) and Common (*Shared*) effects for all predictor variables as proportion of the regression effect (i.e. multiple R^2^) for MR models with and without distance as additional regressor and for the three FC metrics: Coherence (**A**), Power Correlations (**B**), Granger Causality (**C**). Common (*Shared*) effects are summed separately for positive and negative effects. Inset displays summed Common negative effects for all pairs of predictors involving beta-band synchronizations, Sum ± SD.

**Figure S8.**
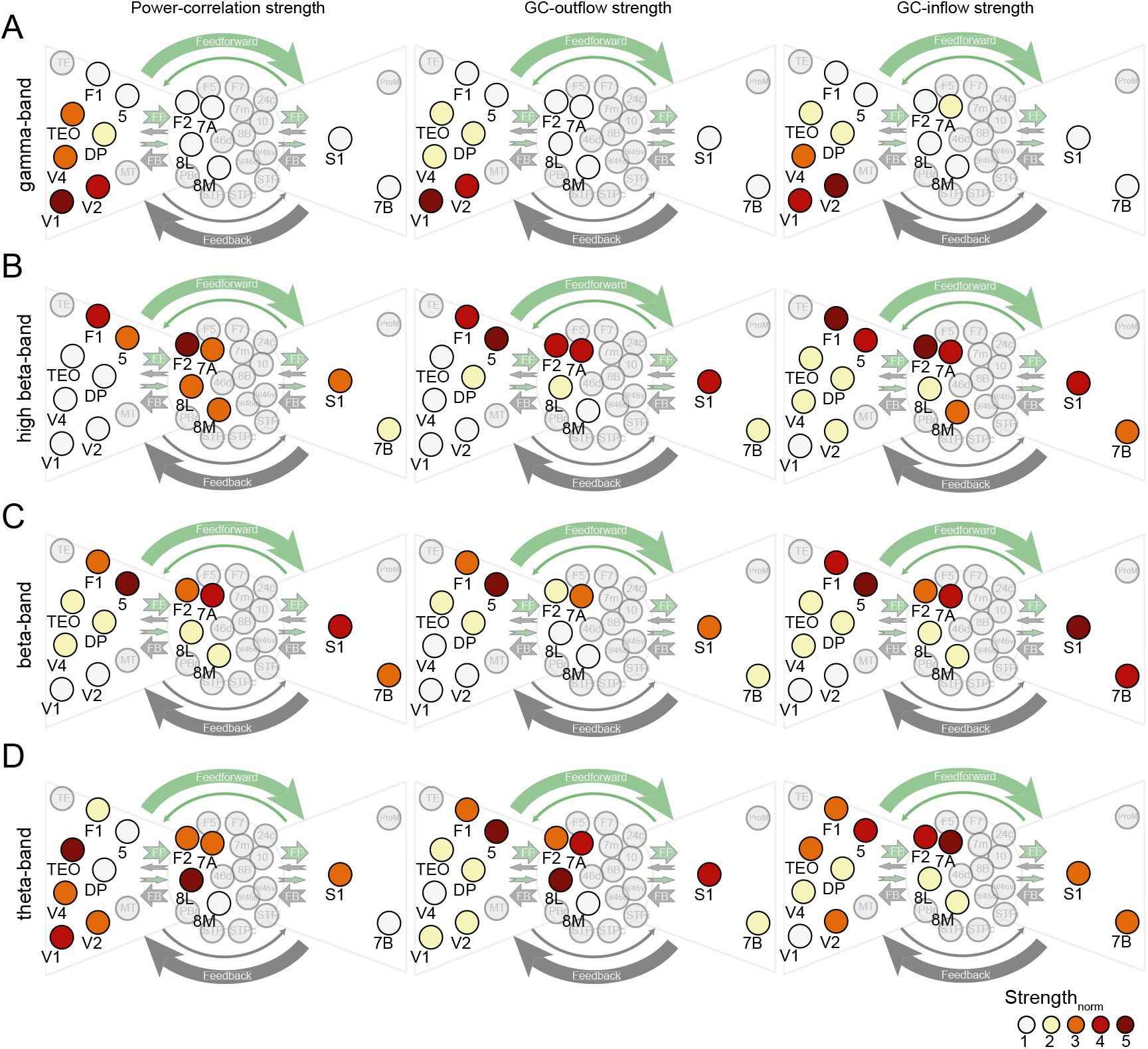
Functional strength integration in the anatomical core-periphery structure. Same as Figure 8, but for power-correlation (left), GC-outflow (middle) and GC-inflow (right) strengths.

**TableS1.** Summary statistics of multiple regression models for FLNe. Relative importance measures (Commonality Analysis: Direct and Shared coefficients, General Dominance – GenDom., adjusted Effect Size and Relative Importance Weights – RIW) performed on the mean over all trials; otherwise, mean ± 99.9% CI estimated from bootstrap over trials. Mean over all trials ± 99.9% CI estimated from bootstrap over trials.

| Predictors | R<sup>2</sup> | R<sub>adj</sub><sup>2</sup> | NRMSE | r<sub>s</sub><sup>2</sup> | Beta values | Beta weights | sig. | Direct | Shared | GenDom. | Effect size | RIW |
| --- | --- | --- | --- | --- | --- | --- | --- | --- | --- | --- | --- | --- |
| | 0.563<br>± 0.02 | 0.522<br>± 0.02 | 0.23<br>± 0.002 | | | | | | | | | |
| Theta | | | | 0.83 ±<br>0.08 | 1.73 ±<br>1.65 | 0.20 ±<br>0.19 | ns | 0.0068 | 0.4606 | 0.153 | 0.136 | 0.138 |
| Beta | | | | 0.34 ±<br>0.02 | 1.33 ±<br>0.26 | 0.28 ±<br>0.05 | < 0.1 | 0.0241 | 0.1914 | 0.066 | 0.105 | 0.079 |
| High-beta | | | | 0.69 ±<br>0.03 | 1.66 ±<br>0.75 | 0.34 ±<br>0.15 | ns | 0.0183 | 0.3919 | 0.125 | 0.203 | 0.123 |
| Gamma | | | | 0.51 ±<br>0.04 | 2.23 ±<br>0.53 | 0.43 ±<br>0.10 | 0.0165 | 0.0562 | 0.2851 | 0.123 | 0.213 | 0.144 |
| Surface dist. | | | | 0.65 ±<br>0.02 | 0.01 ±<br>.003 | 0.17 ±<br>0.04 | ns | 0.0056 | 0.3678 | 0.096 | -0.093 | 0.080 |
| | 0.569<br>± 0.01 | 0.528<br>± 0.01 | 0.23<br>± 0.002 | | | | | | | | | |
| Theta | | | | 0.83 ±<br>0.08 | 1.58 ±<br>1.59 | 0.19 ±<br>0.19 | ns | 0.0057 | 0.4618 | 0.144 | 0.123 | 0.133 |
| Beta | | | | 0.34 ±<br>0.02 | 0.76 ±<br>0.22 | 0.16 ±<br>0.04 | ns | 0.0064 | 0.1850 | 0.057 | 0.057 | 0.068 |
| High-beta | | | | 0.69 ±<br>0.03 | 1.04 ±<br>0.76 | 0.21 ±<br>0.15 | ns | 0.0061 | 0.3858 | 0.115 | 0.119 | 0.114 |
| Gamma | | | | 0.50 ±<br>0.04 | 1.52 ±<br>0.57 | 0.29 ±<br>0.11 | ns | 0.0206 | 0.2645 | 0.104 | 0.141 | 0.123 |
| WM dist. | | | | 0.84 ±<br>0.02 | -0.02 ±<br>.005 | -0.21 ±<br>0.04 | ns | 0.0072 | 0.4674 | 0.146 | 0.126 | 0.128 |
| | 0.558<br>± 0.01 | 0.525<br>± 0.02 | 0.23<br>± 0.002 | | | | | | | | | |
| Theta | | | | 0.84 ±<br>0.08 | 1.56 ±<br>1.60 | 0.18 ±<br>0.19 | ns | 0.0055 | 0.4619 | 0.184 | 0.121 | 0.162 |
| Beta | | | | 0.34 ±<br>0.02 | -1.14 ±<br>0.22 | 0.24 ±<br>0.04 | < 0.1 | 0.0188 | 0.1726 | 0.080 | 0.086 | 0.091 |
| High-beta | | | | 0.70 ±<br>0.03 | 1.44 ±<br>0.75 | 0.29 ±<br>0.15 | ns | 0.0137 | 0.3782 | 0.152 | 0.166 | 0.146 |
| Gamma | | | | 0.51 ±<br>0.04 | 2.02 ±<br>0.53 | 0.39 ±<br>0.10 | 0.02 | 0.0511 | 0.2340 | 0.142 | 0.186 | 0.160 |

**Table S1 (continued). Summary statistics of multiple regression models for FLNe.** 81 Mean over all trials ± 99.9% CI estimated from bootstrap over trials. 83 Power correlations
| Predictors | R<sup>2</sup> | R<sub>adj</sub><sup>2</sup> | NRMSE | r<sub>s</sub><sup>2</sup> | Beta values | Beta weights | sig. | Direct | Shared | GenDom. | Effect size | RIW |
| --- | --- | --- | --- | --- | --- | --- | --- | --- | --- | --- | --- | --- |
| | 0.545<br>± 0.08 | 0.502<br>± 0.08 | 0.23<br>± 0.01 | | | | | | | | | |
| Theta | | | | 0.43<br>± 0.28 | 0.83<br>± 1.21 | 0.19<br>± 0.24 | ns | 0.0164 | 0.2193 | 0.092 | 0.086 | 0.094 |
| Beta | | | | 0.25<br>± 0.07 | 2.03<br>± 0.75 | 0.49<br>± 0.18 | 0.0028 | 0.0859 | 0.0525 | 0.107 | 0.167 | 0.118 |
| High-beta | | | | 0.29<br>± 0.18 | 0.05<br>± 0.86 | 0.01<br>± 0.20 | ns | 0.0001 | 0.1565 | 0.051 | 0.004 | 0.049 |
| Gamma | | | | 0.39<br>± 0.14 | 2.44<br>± 1.06 | 0.49<br>± 0.21 | 0.0011 | 0.1051 | 0.1103 | 0.149 | 0.210 | 0.159 |
| Surface dist. | | | | 0.67<br>± 0.10 | -0.01<br>± 0.01 | -0.14<br>± 0.18 | ns | 0.0070 | 0.3608 | 0.146 | 0.079 | 0.125 |
| | 0.571<br>± 0.05 | 0.529<br>± 0.06 | 0.23<br>± 0.007 | | | | | | | | | |
| Theta | | | | 0.41 ± 0.26 | 0.61 ± 1.15 | 0.14<br>± 0.23 | ns | 0.0087 | 0.2770 | 0.086 | 0.063 | 0.089 |
| Beta | | | | 0.24 ± 0.06 | -1.50 ± 0.77 | 0.36<br>± 0.18 | 0.0316 | 0.043 | 0.0981 | 0.089 | 0.124 | 0.100 |
| High-beta | | | | 0.27 ± 0.17 | 0.11 ± 0.73 | 0.03<br>± 0.17 | ns | 0.0004 | 0.1562 | 0.050 | 0.010 | 0.049 |
| Gamma | | | | 0.38 ± 0.14 | 1.97<br>± 1.00 | 0.40<br>± 0.20 | 0.0085 | 0.0618 | 0.1536 | 0.129 | 0.170 | 0.139 |
| WM dist. | | | | 0.83 ± 0.08 | -0.03 ± 0.02 | -0.32<br>± 0.17 | < 0.1 | 0.0321 | 0.4425 | 0.216 | 0.204 | 0.194 |
| | 0.539<br>± 0.10 | 0.504<br>± 0.11 | 0.23<br>± 0.01 | | | | | | | | | |
| Theta | | | | 0.39 ± 0.26 | 1.25 ± 1.32 | 0.24<br>± 0.26 | < 0.1 | 0.0281 | 0.2076 | 0.125 | 0.105 | 0.122 |
| Beta | | | | 0.27 ± 0.08 | 2.34 ± 0.71 | 0.56<br>± 0.16 | 0.0001 | 0.1516 | -0.0132 | 0.151 | 0.191 | 0.155 |
| High-beta | | | | 0.31 ± 0.19 | 0.15 ± 0.94 | 0.03<br>± 0.22 | ns | 0.0003 | 0.1563 | 0.000 | 0.010 | 0.062 |
| Gamma | | | | 0.36 ± 0.14 | 2.70 ± 1.11 | 0.55<br>± 0.21 | 0.0001 | 0.1574 | 0.0580 | 0.157 | 0.233 | 0.199 |

| Predictors | $R^2$ | $R_{adj}^2$ | NRMSE | $r_s^2$ | Beta values | Beta weights | sig. | Direct | Shared | GenDom. | Effect size | RIW |
| --- | --- | --- | --- | --- | --- | --- | --- | --- | --- | --- | --- | --- |
| | 0.485<br>± 0.02 | 0.458<br>± 0.02 | 0.23<br>± 0.02 | | | | | | | | | |
| Theta | | | | 0.61 ± 0.07 | 0.24 ± 0.27 | 0.07 ± 0.07 | ns | 0.0009 | 0.2963 | 0.079 | 0.037 | 0.073 |
| Beta | | | | 0.29 ± 0.03 | -0.09 ± 0.17 | -0.04 ± 0.07 | ns | 0.0002 | 0.1412 | 0.039 | -0.014 | 0.039 |
| High-beta | | | | 0.68 ± 0.03 | 0.78 ± 0.20 | 0.36 ± 0.08 | < 0.1 | 0.0162 | 0.3162 | 0.100 | 0.191 | 0.090 |
| Gamma | | | | 0.87 ± 0.03 | 1.17 ± 0.17 | 0.53 ± 0.06 | 0.0004 | 0.0751 | 0.3465 | 0.183 | 0.316 | 0.199 |
| Surface dist. | | | | 0.67 ± 0.02 | -0.01 ± 0.003 | 0.09 ± 0.05 | ns | 0.0016 | 0.3220 | 0.085 | -0.045 | 0.084 |
| | 0.492<br>± 0.01 | 0.465<br>± 0.01 | 0.23<br>± 0.001 | | | | | | | | | |
| Theta | | | | 0.60 ± 0.07 | 0.19 ± 0.29 | 0.06 ± 0.08 | ns | 0.0005 | 0.2967 | 0.074 | 0.030 | 0.069 |
| Beta | | | | 0.29 ± 0.02 | -0.08 ± 0.17 | -0.03 ± 0.07 | ns | 0.0002 | 0.1414 | 0.037 | -0.012 | 0.036 |
| High-beta | | | | 0.67 ± 0.03 | 0.44 ± 0.16 | 0.20 ± 0.06 | ns | 0.0052 | 0.3324 | 0.092 | 0.108 | 0.083 |
| Gamma | | | | 0.86 ± 0.03 | 0.89 ± 0.17 | 0.40 ± 0.06 | 0.007 | 0.0414 | 0.4216 | 0.160 | 0.240 | 0.178 |
| WM dist. | | | | 0.85 ± 0.02 | -0.02 ± 0.004 | -0.21 ± 0.04 | ns | 0.0086 | 0.4202 | 0.130 | 0.127 | 0.126 |
| | 0.484<br>± 0.02 | 0.462<br>± 0.02 | 0.23<br>± 0.002 | | | | | | | | | |
| Theta | | | | 0.61 ± 0.07 | 0.24 ± 0.27 | 0.07 ± 0.07 | ns | 0.0009 | 0.2963 | 0.097 | 0.038 | 0.087 |
| Beta | | | | 0.29 ± 0.03 | -0.09 ± 0.17 | -0.04 ± 0.07 | ns | 0.0002 | 0.1412 | 0.048 | -0.014 | 0.045 |
| High-beta | | | | 0.69 ± 0.03 | 0.68 ± 0.19 | 0.31 ± 0.07 | ns | 0.0148 | 0.3176 | 0.125 | 0.165 | 0.111 |
| Gamma | | | | 0.87 ± 0.03 | 1.09 ± 0.15 | 0.49 ± 0.05 | 0.0002 | 0.0831 | 0.3385 | 0.214 | 0.295 | 0.240 |

**TableS2.** Exponential decay rates of FC measures with interareal distance. Mean over all trials ± SEM estimated from bootstrap over trials.

|  | Interareal distance | Coherence | Power Correlations | Granger Causality |
| --- | --- | --- | --- | --- |
| Theta | WM | 0.024 ± 0.0005 mm<sup>-1</sup> | 0.028 ± 0.004 mm<sup>-1</sup> | 0.055 ± 0.0012 mm<sup>-1</sup> |
| | surface | 0.016 ± 0.004 mm<sup>-1</sup> | 0.018 ± 0.003 mm<sup>-1</sup> | 0.034 ± 0.0008 mm<sup>-1</sup> |
| Beta | WM | 0.033 ± 0.0003 mm<sup>-1</sup> | 0.032 ± 0.001 mm<sup>-1</sup> | 0.063 ± 0.0007 mm<sup>-1</sup> |
| | surface | 0.021 ± 0.0002 mm<sup>-1</sup> | 0.019 ± 0.001 mm<sup>-1</sup> | 0.039 ± 0.0005 mm<sup>-1</sup> |
| High-Beta | WM | 0.042 ± 0.0004 mm<sup>-1</sup> | 0.028 ± 0.003 mm<sup>-1</sup> | 0.085 ± 0.0008 mm<sup>-1</sup> |
| | surface | 0.027 ± 0.0003 mm<sup>-1</sup> | 0.018 ± 0.002 mm<sup>-1</sup> | 0.055 ± 0.0005 mm<sup>-1</sup> |
| Gamma | WM | 0.027 ± 0.0004 mm<sup>-1</sup> | 0.015 ± 0.002 mm<sup>-1</sup> | 0.075 ± 0.0008 mm<sup>-1</sup> |
| | surface | 0.018 ± 0.0003 mm<sup>-1</sup> | 0.010 ± 0.001 mm<sup>-1</sup> | 0.049 ± 0.0006 mm<sup>-1</sup> |

## Notes

### Summary of Updates

Original version of the submitted manuscript.

